# PIE-1 promotes SUMOylation and activation of HDAC1 during the *C. elegans* oogenesis

**DOI:** 10.1101/2020.08.17.253955

**Authors:** Heesun Kim, Yue-He Ding, Shan Lu, Mei-Qing Zuo, Darryl Conte, Meng-Qiu Dong, Craig C. Mello

**Author notes:** These authors contributed equally (Heesun Kim, Yue-He Ding). Corresponding author Craig C. Mello, RNA Therapeutics Institute, University of Massachusetts Medical School, 368 Plantation Street, AS5-2049, Worcester, MA 01605, USA.

## Abstract

Germlines shape and balance heredity, integrating and regulating information from both parental and foreign sources. Insights into how the germline handles information have come from the identification of factors that specify or maintain the germline fate. In early *C. elegans* embryos, the CCCH zinc-finger protein PIE-1 localizes to the germline where it prevents somatic differentiation programs. Here we show that PIE-1 also functions in the meiotic ovary where it becomes SUMOylated and engages the SUMO-conjugating machinery. Using whole-proteome mass spectrometry to detect SUMO-conjugated proteins, we identify HDAC SUMOylation as a target of PIE-1. Our findings suggest that SUMOylation activates HDAC, lowering histone acetylation and enhancing Argonaute-mediated surveillance in the germline.

## Introduction

During every life cycle the eukaryotic germline orchestrates a remarkable set of informational tasks that shape heredity and create variation necessary for the evolution of new species. One approach for understanding the mechanisms that promote germline specification and function has been the identification of genes whose protein products localize exclusively to the germline and for which loss of function mutations result in absent or non-functional germ cells and gametes (Seydoux and Braun, 2006). In *C. elegans*, PIE-1 is a key regulator of germline specification (Mello et al., 1992). The C. elegans zygote, P0, undergoes a series of asymmetric divisions that generate 4 somatic founder cells and the germline blastomere P4. The PIE-1 protein is maternally deposited and uniformly present in the cytoplasm and nucleus of the zygote, but rapidly disappears in each somatic blastomere shortly after division (Mello et al., 1996; Reese et al., 2000; Tenenhaus et al., 1998). In *pie-1* mutants, the germline lineage differentiates into extra intestinal cells causing an embryonic arrest (Mello et al., 1992). PIE-1 localizes prominently in nuclei in the early P-lineage blastomeres, and persists in the primordial embryonic germline through much of embryogenesis (Mello et al., 1996). The presence of PIE-1 correlates with global hypo-phosphorylation of the C-terminal domain (CTD) of RNA polymerase II (pol-II) in germline blastomeres (Seydoux and Dunn, 1997), and some studies suggest that PIE-1 may directly inhibit the CTD kinase to prevent transcriptional activation (Batchelder et al., 1999; Ghosh and Seydoux, 2008).

PIE-1 is a member of the tandem CCCH Zinc finger protein family (Blackshear P.J., 2005). PIE-1 differs from most of its homologs in having a prominent nuclear localization. However, like most of its family members, PIE-1 also localizes in the cytoplasm where it is thought to bind and regulate the expression of germline mRNAs including the *nos-2* mRNA (Tenenhaus et al., 2001). Hints at the nuclear function of PIE-1 came from a yeast two-hybrid screen which identified the Krüppel-type zinc-finger protein MEP-1 as a PIE-1 interacting factor (Unhavaithaya et al., 2002). MEP-1 co-purifies with LET-418, a homolog of mammalian ATP-dependent nucleosome remodeling factor Mi-2 (von Zelewsky et al., 2000), and with HDA-1, a homolog of mammalian histone deacetylase HDAC1 (Shi and Mello, 1998). Inactivation of maternal *mep-1* and *let-418* causes a striking developmental arrest of L1-stage larvae, whose somatic cells adopt germline-specific transcriptional programs, and assemble germline-specific peri-nuclear nuage-like structures called P-granules (Unhavaithaya et al., 2002). The soma-to-germline transformation and developmental arrest phenotypes depend on the trithorax-related protein MES-4, and components of a polycomb repressive complex (PRC2) (MES-2 and MES-3) (Unhavaithaya et al., 2002), whose functions are thought to promote fertility by maintaining germline chromatin (Strome and Updike, 2015). Taken together, these previous studies on PIE-1 suggest that it functions as a master-regulator of the germline fate in *C. elegans* embryos, preventing somatic differentiation, while also protecting the germline chromatin from remodeling. However, the possible biochemical mechanisms through which this small CCCH-zinc finger protein exerts its dual effects on transcription and chromatin in the germline were entirely unknown.

Here we show PIE-1 promotes the regulation of its targets at least in part through the small ubiquitin-like modifier (SUMO). By yeast two-hybrid screening we show that PIE-1 engages the highly conserved E2 SUMO ligase UBC-9. UBC-9 enzymes catalyze the addition of the small ubiquitin-like modifier (SUMO) to lysine residues on target proteins (Capili and Lima, 2007; Geiss-Friedlander and Melchior, 2007; Johnson, 2004). The reversible addition of SUMO (or SUMOylation) and its removal by de-SUMOylating enzymes is thought to occur on thousands of substrate proteins with diverse functions, especially nuclear functions including DNA replication, chromatin silencing, and the DNA damage response (Hendriks and Vertegaal, 2016). SUMOylation can have multiple effects, and is not primarily associated with the turnover of its targets, but rather is often associated with changes in protein interactions. For example, SUMOylation of a protein can promote interactions with proteins that contain SUMO-interacting motifs (SIMs) (Matunis et al., 2006; Psakhye and Jentsch, 2012; Shen et al., 2006).

Through a series of genetic and biochemical studies, we show that the SUMO pathway promotes the activity of PIE-1 in preserving the embryonic germline. We show that PIE-1 is itself modified by SUMO on lysine 68. Paradoxically, PIE-1 is not SUMOylated in early embryos, but rather in adult animals where PIE-1 was not previously known to be expressed or functional. Indeed, CRISPR-mediated GFP tagging of the endogenous *pie-1* locus confirmed uniform nuclear expression of PIE-1 protein throughout the meiotic zone and in oocytes of adult hermaphrodites. Using whole proteome analysis for detecting SUMO-conjugated proteins, we identify the type 1 HDAC, HDA-1, as a protein modified by SUMO in a PIE-1-dependent manner. Surprisingly, whereas PIE-1 was originally thought to inhibit the MEP-1/Mi-2/HDA-1 complex in embryos, we show that in the germline PIE-1 acts in concert with the SUMO pathway to promote the association of MEP-1 with HDA-1, and to maintain the hypoacetylation of germline chromatin. Our findings along with those from a parallel study are consistent with a model in which the assembly of a MEP-1/Mi-2/HDA-1 chromatin remodeling complex is tightly regulated in the hermaphrodite germline to coordinate Argonaute-mediated genome surveillance.

## Results

### PIE-1 is SUMOylated on K68 in adult germ cells

To explore how PIE-1 promotes germline specification we sought to identify protein interactors. Purification of PIE-1 protein from embryo extracts proved to be challenging because the protein is expressed transiently in early embryos, where it is only present in early germline cells. Moreover, PIE-1 was insoluble and unstable in worm lysates, preventing the analysis of PIE-1 complexes by immunoprecipitation (Figures S1A-S1C). We therefore performed a yeast two-hybrid screen to identify PIE-1 interactors (Figure 1A and Table S1; see Materials and Methods). As expected, this screen identified the Krüppel-type zinc finger protein MEP-1, a known PIE-1 interactor and co-factor of the Mi-2/NuRD (nucleosome remodeling deacetylase) complex (Passannante et al., 2010; Unhavaithaya et al., 2002). In addition, we identified the small ubiquitin-like modifier SMO-1 (SUMO); the E2 SUMO-conjugating enzyme UBC-9 (Jones et al., 2002); and GEI-17 a homolog of Drosophila Su(var)2-10 an E3 SUMO ligase (Holway et al., 2005).

**Figure 1.**
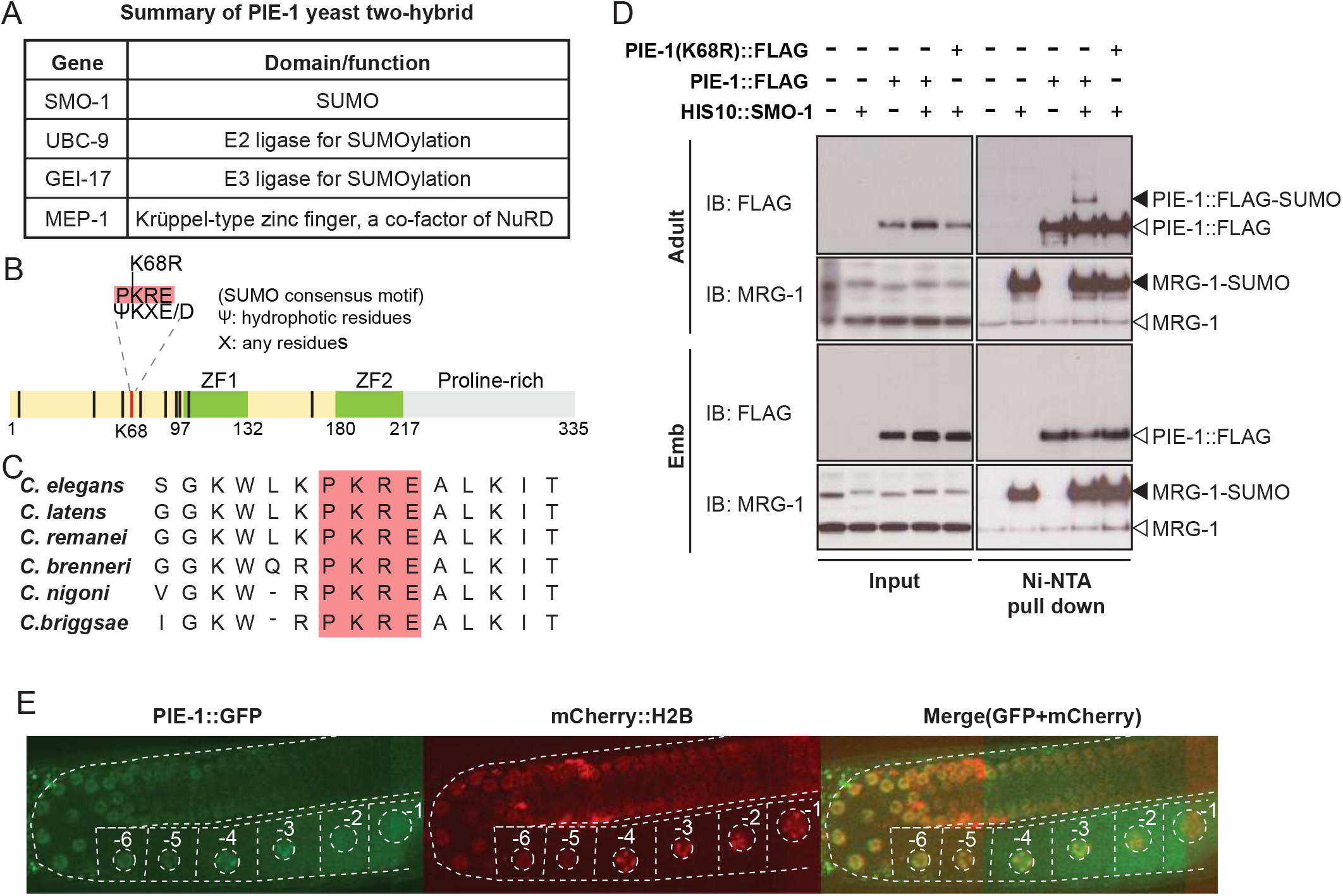
PIE-1 is SUMOylated on K68 residues in the *C. elegans* germline. (A) Summary of PIE-1 interactors identified in yeast two-hybrid screen (see Table S1 for all identified PIE-1 interactors). (B) Schematic of PIE-1 containing the consensus SUMO acceptor site (ΨKXE, where Ψ represents a hydrophobic amino acid and X is any amino acids), K68. The black bars represent lysine residues on PIE-1 and the K68 is indicated with red bar. (C) Partial sequence alignment of the consensus SUMO acceptor site (red box) of PIE-1 over other sister organisms. (D) *in vivo* SUMO purification assay followed by western blotting in adult and early embryo lysates. The black triangles represent SUMOylated PIE-1 and MRG-1. Unmodified PIE-1 and MRG-1 are indicated by the white triangles. (E) Confocal live images of adult germline expressing PIE-1::GFP and mCherry::H2B in *pie-1::gfp; pie-1p::mCherry::his-58* strain.

Motif analysis predicted one consensus SUMO acceptor site (ΨKXE; Rodriguez et al., 2001) in PIE-1 that is perfectly conserved in PIE-1 orthologs of other *Caenorhabditis* species (Figures 1B and 1C). To test if SUMO is conjugated to PIE-1, we inserted a poly-histidine epitope into the endogenous *smo-1* gene, and then used nickel affinity chromatography to enrich SUMOylated proteins from worm lysates (Tatham et al., 2009). We found that 10 histidines at the N-terminus of SUMO allowed good retention of SUMO and SUMO-conjugated proteins to the nickel affinity column under stringent denaturing and washing conditions (Figures S2A-S2C). Mass spectrometry of eluates from the nickel affinity column revealed enrichment of SUMO and 990 other proteins from *his10::smo-1* lysates compared to control lysates from untagged *smo-1* worms or *smo-1(RNAi)* worms (Table S2). SUMOylated MRG-1, for example, was strongly enriched by nickel affinity chromatography of *his10::smo-1* lysates, as detected by mass spectrometry and western blot analyses (Table S2 and Figure 1D).

PIE-1 is not an abundant protein and was not detected by mass spectrometry of SUMOylated proteins enriched by nickel affinity chromatography. To test if western blot analysis would be a more sensitive approach, we inserted a sequence encoding the FLAG epitope into the endogenous *pie-1* gene, generating a homozygous viable *pie-1::flag* allele. The PIE-1::FLAG protein was expressed at similar levels in WT and *his10::smo-1* animals (Figure 1D). Nickel affinity chromatography enriched a modified form of PIE-1::FLAG that depended on expression of HIS10::SMO-1 (Figure 1D). Importantly, the modified PIE-1::FLAG protein was ~15 kD larger than unmodified PIE-1::FLAG and was not detected when we mutated the presumptive SUMO acceptor site lysine 68 to arginine in PIE-1 (K68R; Figure 1D). These data suggest that PIE-1 is SUMOylated on lysine 68.

We were surprised to detect SUMOylated PIE-1 in adult hermaphrodites but not in embryos (Figure 1D). Previous studies had only detected PIE-1 protein within embryonic germ cells and proximal oocytes of adult hermaphrodites (Reese et al., 2000; Tenenhaus et al., 1998). We therefore carefully monitored PIE-1::GFP robustly expressed from the endogenous *pie-1* locus (Kim et al., 2014). As expected, in embryonic germ cells PIE-1::GFP localized to nuclei, cytoplasm, and cytoplasmic P-granules (i.e., P lineage; Figure S3) (Mello et al., 1996; Reese et al., 2000; Tenenhaus et al., 1998). In the adult germline, we found that PIE-1::GFP colocalized with chromosomes in germ nuclei beginning at the pachytene stage, with levels gradually increasing during oocyte maturation (Figures 1E and S3). These findings suggest that SUMOylation of PIE-1 occurs within a heretofore unexplored zone of PIE-1 expression within the maternal germline.

### PIE-1 SUMOylation and SUMO pathway factors function together to promote fertility and embryonic development

To investigate the functional significance of PIE-1 SUMOylation, we tested for genetic interactions between *pie-1* and SUMO pathway mutants. In early embryos, maternal PIE-1 promotes the germ cell fate of the P2 blastomere and prevents P2 from adopting the endoderm and mesoderm fates of its somatic sister blastomere, EMS (Mello et al., 1992). Hermaphrodite worms homozygous for the loss-of-function *pie-1(zu154)* allele are fertile, but 100% of their embryos arrest development with extra pharyngeal and intestinal cells (Mello et al., 1992). By contrast, we found that homozygous *pie-1(ne4303)* hermaphrodites expressing PIE-1(K68R) produced mostly viable and fertile progeny, but showed variable fertility phenotype which resulted in significantly reduced fertility and increased embryonic lethality (Figures 2A and 2B), suggesting that the K68R lesion causes a partial loss of PIE-1 function.

**Figure 2.**
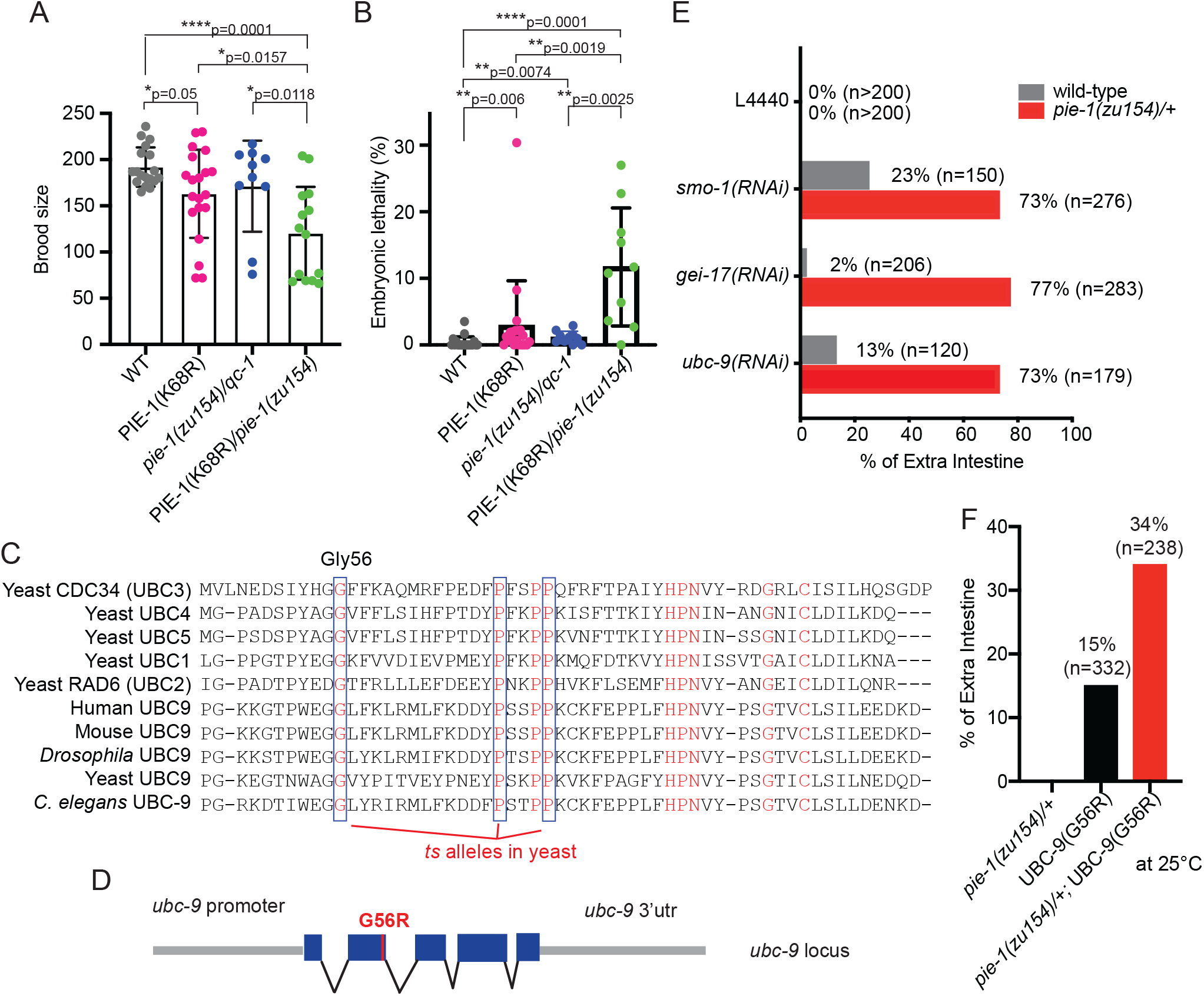
Genetic interactions between *pie-1* and SUMO pathway. (A) Brood sizes and (B) embryonic lethality of wild-type (N2), *pie-1(ne4303[K68R]), pie-1(zu154)/qc-1, and pie-1(ne4303[K68R])/pie-1(zu154)*. Wilcoxon-Mann-Whitney: *P≤0.05, **P≤0.01, ****P≤0.0001 (C) Partial sequence alignment of UBC enzymes including UBC9. Residues conserved in all UBC proteins are shown in red. The indicated G56, P69, and P73 residues are isolated *ts* alleles in yeast Cdc34 and both are conserved in all UBC proteins over other organisms (blue box). Among many conserved residues, the G56 residue is identified as a *ts* allele in *C. elegans.* (D) Schematic of *ubc-9* genomic locus showing G56R position. (E) Genetic interaction of *pie-1* with SUMO pathway. Defective SUMOylation pathway by RNAi feeding-mediated each indicated gene knockdown causes high frequency of *pie-1* dead embryos at 20°C. (F) Genetic interaction of *pie-1* with *ubc-9(ne4446[G56R]) ts* allele. The *ubc-9(ne4446[G56R]) ts* allele makes 2-fold increased *pie-1* dead embryos at 25°C, but shows a lower penetrance extra gut derived from the germline blastomere compared to RNAi feeding experiments.

We found that further lowering *pie-1* activity by placing *pie-1(ne4303[K68R])* over a null allele, *pie-1(zu154),* dramatically enhanced deficits in fertility (Figure 2A) and in embryo viability (Figure 2B). For example, about 11% of embryos produced by trans-heterozygotes failed to hatch (Figure 2B). The production of excess intestinal cells, a hallmark of *pie-1* loss of function mutants, is readily scored by light microcopy (Mello et al., 1992). We therefore carefully scored a subset of the arrested embryos produced by *pie-1(zu154)* trans-heterozygotes and found that 64% (14/22) embryos examined exhibited supernumerary intestinal cells consistent with a *pie-1* loss-of-function phenotype. Thus the PIE-1(K68R) substitution causes a partial loss of PIE-1 function.

Null alleles of *smo-1* and *ubc-9* cause recessive sterile phenotypes (Broday et al., 2004; Lim et al., 2014; Reichman et al., 2018), suggesting that their zygotic expression is required for germline development and function. Null alleles of gei-17, in contrast, were homozygous viable but exhibited reduced hermaphrodite fertility and embryo viability (Kim et al., parallel). In order to address whether these factors contribute to PIE-1-dependent embryo viability, we sought to deplete the function of SUMO-pathway genes by RNAi beginning at the L4 larval stage, after sperm production and the transition to oogenesis were completed. We also used genome editing to engineer a temperature-sensitive *ubc-9(ne4446)* allele expressing a G56R amino-acid substitution modeled after temperature-sensitive alleles of related yeast enzymes (Figures 2C and 2D)(Betting and Seufert, 1996; Prendergast et al., 1995). Depletion of GEI-17 by RNAi at the L4 stage caused embryo lethality among ~19% (39/206) of progeny scored and 10% of these arrested embryos (4/39) exhibited extra intestinal cells (Figure 2E). RNAi of *smo-1* and *ubc-9*, or shifting *ubc-9(ne4446[G56R])* mutants to non-permissive temperature (25°C) beginning at the L4 stage, resulted in fertile adults that produced 100% dead embryos (n=150, n=120, and n=332, respectively). The embryos arrested development with defective morphology but well-differentiated tissues. Moreover, 23% of *smo-1(RNAi)*, 13% of *ubc-9(RNAi)*, and 15% of *ubc-9(ne4446[G56R])* mutant embryos arrested development with extra intestinal cells (Figures 2E and 2F). These findings are consistent with previous cell lineage studies on *smo-1(RNAi)* and *ubc-9(RNAi)* embryos that demonstrated the P2 blastomere expresses EMS-like cell lineage patterns, a *pie-1* mutant phenotype (Santella et al., 2016). Taken together with our examination of intestinal cell numbers these findings suggest that SUMO-pathway mutants exhibit *pie-1*-like transformations in embryonic cell fate.

If, as suggested by the above findings, SUMO functions along with PIE-1, we reasoned that further reducing the *pie-1* gene dose should increase the frequency of *smo-1*, *ubc-9*, or *gei-17* dead embryos with *pie-1-*like embryonic phenotypes. Consistent with this idea we found that depletion of the SUMO factors in L4-stage *pie-1(zu154)* heterozygotes (which by themselves produce fully viable progeny) resulted in enhanced frequencies of embryos with excess intestinal cells (Figures 2E and 2F). The synergy between *pie-1* and *gei-17* was particularly striking: whereas *gei-17(RNAi)* in otherwise wild-type worms caused only a moderate level of embryo lethality, *gei-17(RNAi)* caused heterozygous *pie-1*/+ worms to make 82% (232/283) dead embryos, of which 94% (218/232) made extra intestinal cells (Figure 2E). Taken together, these data suggest that the SUMO pathway promotes PIE-1 function.

### PIE-1 SUMOylation promotes HDA-1 SUMOylation in the germline

SUMO is known to modulate the interactions and activities of target proteins (Geiss-Friedlander and Melchior, 2007; Gill, 2004). We therefore wondered if PIE-1 recruits the SUMO machinery in order to covalently modify and regulate its downstream targets. A previous study suggested that PIE-1 regulates conserved components of the Nucleosome Remodeling and Deacetylase (NuRD) complex, including MEP-1 and its binding partners LET-418/Mi-2 and HDA-1/HDAC1 (Unhavaithaya et al., 2002). We noticed that both HDA-1 and LET418 were identified as SUMOylated proteins in our SUMO proteomics studies. To explore the possibility that PIE-1 promotes the SUMOylation of these factors, we used nickel affinity chromatography to pull down proteins modified with HIS10::SMO-1 and performed western blots to detect MEP-1 and HDA-1. To monitor SUMOylation of MEP-1, we used CRISPR to insert tandem *gfp* and *3xflag* coding sequences into the *mep-1* gene—i.e., *mep-1::gfp::tev::3xflag* (*mep-1::gtf*). For HDA-1 detection, we used a previously validated HDA-1 specific antibody (Beurton et al., 2019). We observed more slowly migrating isoforms of both MEP-1 and HDA-1 after nickel affinity chromatography from *his10::smo-1* lysates (Figures 3A-3D). These protein isoforms were ~15 kD larger than unmodified MEP-1 and HDA-1 and were absent in lysates prepared from *smo-1(RNAi)* worms (Figure 3A), indicating that the larger isoforms result from SUMOylation.

**Figure 3.**
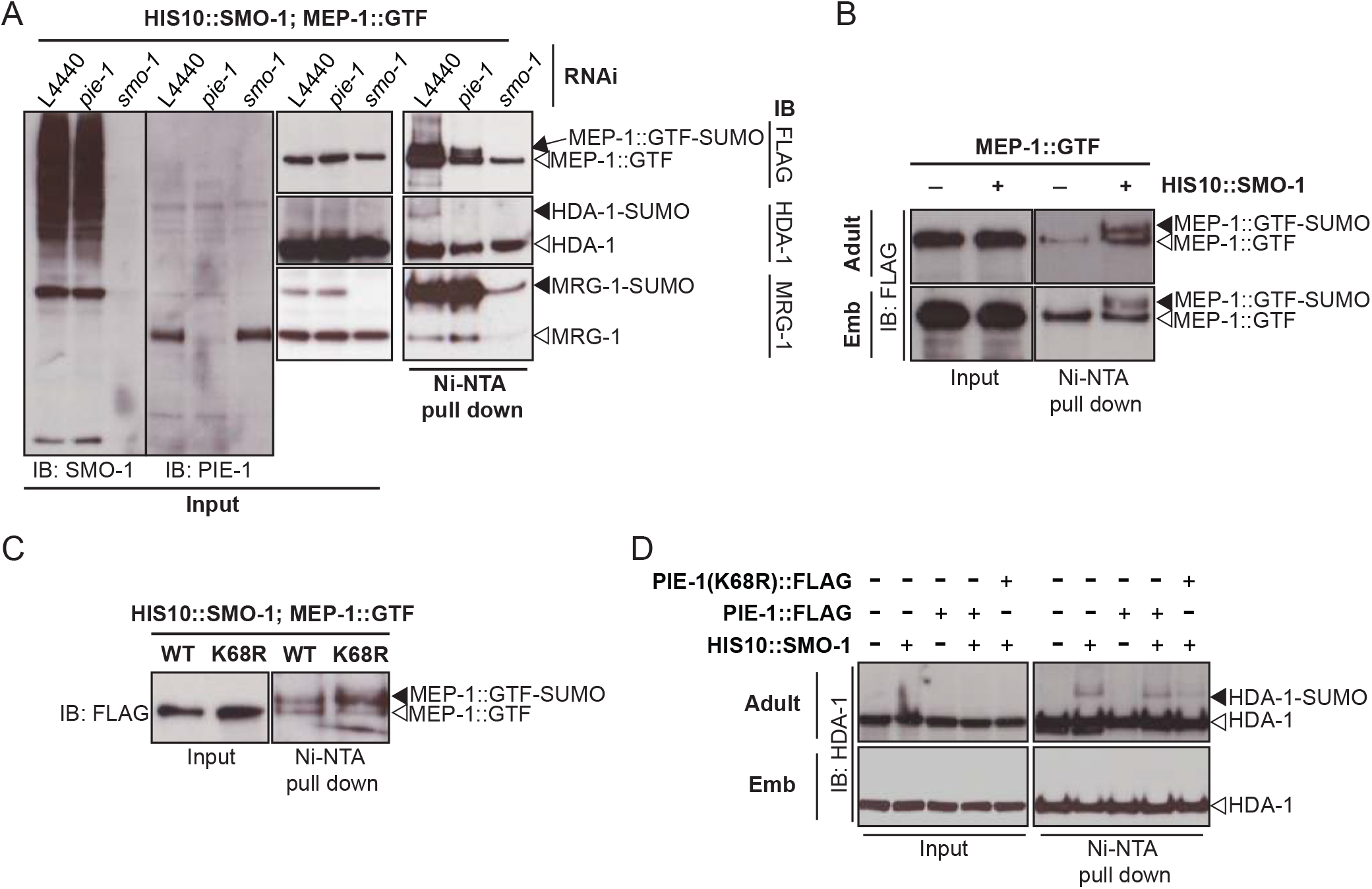
PIE-1 SUMOylation promotes HDA-1 SUMOylation in the adult germline. (A) *in vivo* SUMO purification assay followed by western blotting in adult lysates revealed that PIE-1 is required for HDA-1 SUMOylation. (B), (C) and (D) *in vivo* SUMO purification assay followed by western blotting in adult and embryo lysates. The black triangles indicate SUMOylated proteins and unmodifed proteins are indicated with the white triangles.

Interestingly, we found that SUMOylated HDA-1 but not SUMOylated MEP-1 was strongly depleted in extracts from *pie-1(RNAi)* adults (Figure 3A) and was also significantly reduced in adult extracts from homozygous mutant animals expressing PIE-1(K68R) (Figures 3C and 3D), suggesting that PIE-1 SUMOylation enhances SUMOylation of HDA-1 but is not required for MEP-1 SUMOylation. Indeed, whereas SUMOylated MEP-1 was present in the adult germline and in early embryos (Figure 3B), SUMOylated HDA-1 (like SUMOylated PIE-1) was only detected in the adult germline and not in early embryos (Figure 3D). As expected, MRG-1 SUMOylation was not affected by *pie-1* depletion (Figure 3A).

### PIE-1 SUMOylation promotes formation of a germline MEP-1/HDA-1 complex

To address whether PIE-1 SUMOylation modulates the interaction of HDA-1 with other NuRD complex components, we purified MEP-1::GTF from early embryo or adult lysates using a GBP nanobody (Rothbauer et al., 2008) and detected LET-418 or HDA-1 by western blot. In embryo extracts, where PIE-1 and HDA-1 are not modified by SUMO, MEP-1 interacted robustly with both LET-418 and HDA-1 (Figure 4A). By contrast, in adults MEP-1 interacted robustly with LET-418, but much less prominently with HDA-1 (Figure 4A). This modest association between MEP-1 and HDA-1 in adults required both *smo-1* and *pie-1* activity (Figure 4B). Thus SUMOylation of PIE-1 promotes the interaction of HDA-1 with MEP-1 in the adult germline, but is not required for their much more robust interaction in embryos (Figure 4A). The finding that HDA-1 binding to MEP-1 was more strongly affected by RNAi of *pie-1* or *smo-1* than by the (K68R) substitution (Figure 4B) suggests that, even though it isn't SUMOylated, PIE-1(K68R) promotes HDA-1 SUMOylation and MEP-1 binding in the germline.

**Figure 4.**
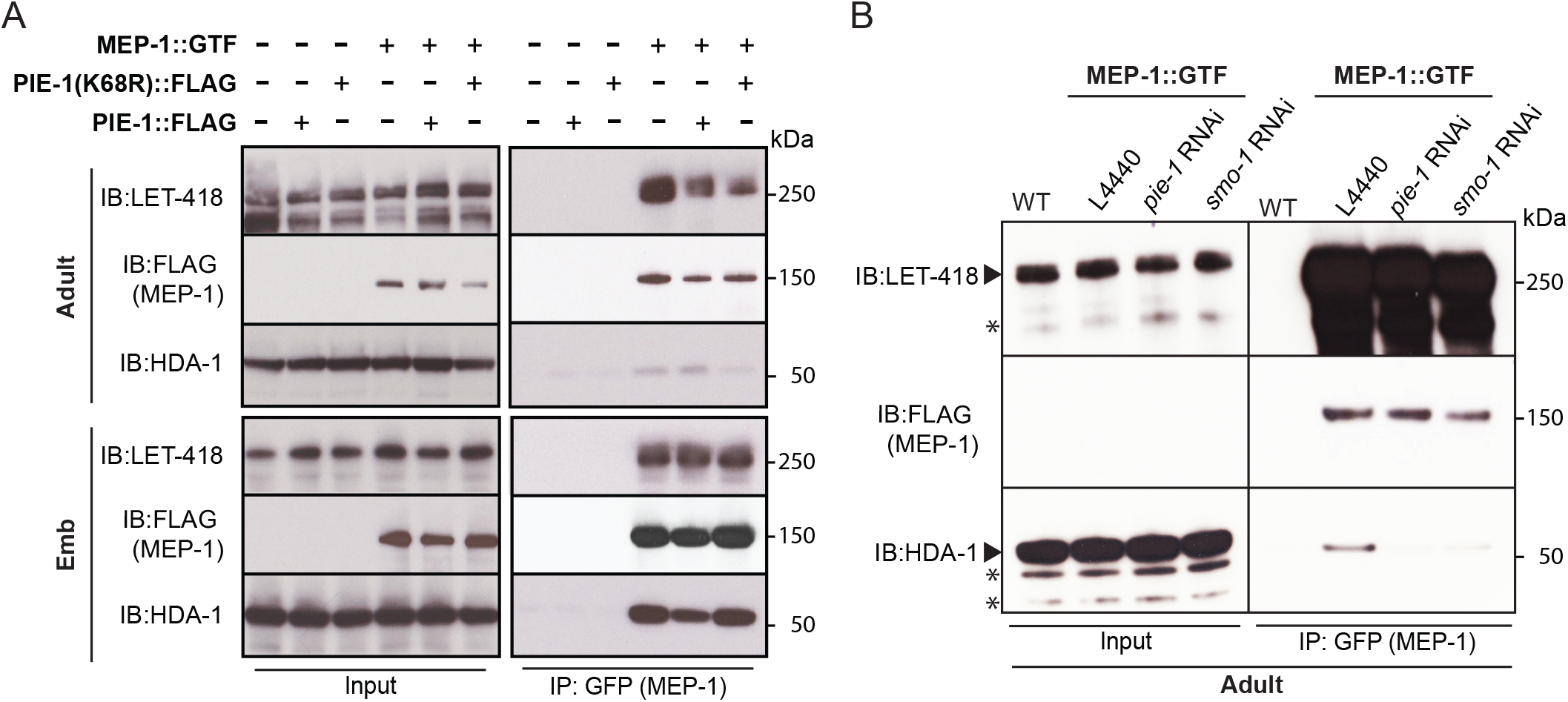
PIE-1 SUMOylation is required for formation of MEP-1/HAD-1 complex in the adult germline. (A) and (B) Co-immunoprecipitation experiment showing physical interaction between MEP-1 and NuRD complex (LET-418 and HDA-1). Immunoprecipitation was performed in embryo and adult lysates using GBP beads, and the indicated proteins (MEP-1, LET-418, and HDA-1) were detected with anti-FLAG antibody, anti-LET-418 antibody, and anti-HDA-1 antibody, respectively. Non-specific bands are indicated with the star.

### PIE-1 suppresses histone acetylation and germline gene expression

If PIE-1 promotes HDA-1 deacetylase activity in the germline, we would expect increased levels of histone acetylation in *pie-1* mutants. Indeed, histone H3 lysine 9 acetylation (H3K9Ac) immunostaining revealed increased levels of H3K9Ac in gonads of homozygous mutant animals expressing PIE-1(K68R) compared to wild-type gonads (Figure 5A), especially in the distal region and in oocytes. To more strongly deplete PIE-1 protein, we engineered an in-frame auxin-responsive PIE-1::DEGRON::GFP (See Materials and Methods). Exposing these animals to auxin from the L1 stage abolished GFP fluorescence in the adult germline (Figure S4) and caused a penetrant *pie-1* maternal-effect embryonic lethal phenotype. Moreover, these auxin treated animals exhibited uniformly high levels of H3K9Ac throughout the germline, at levels higher than those observed in PIE-1(K68R) homozygotes (Figure 5A). These findings suggest that SUMOylation of PIE-1 on K68 enhances its ability to activate HDA-1 in the adult germline.

**Figure 5.**
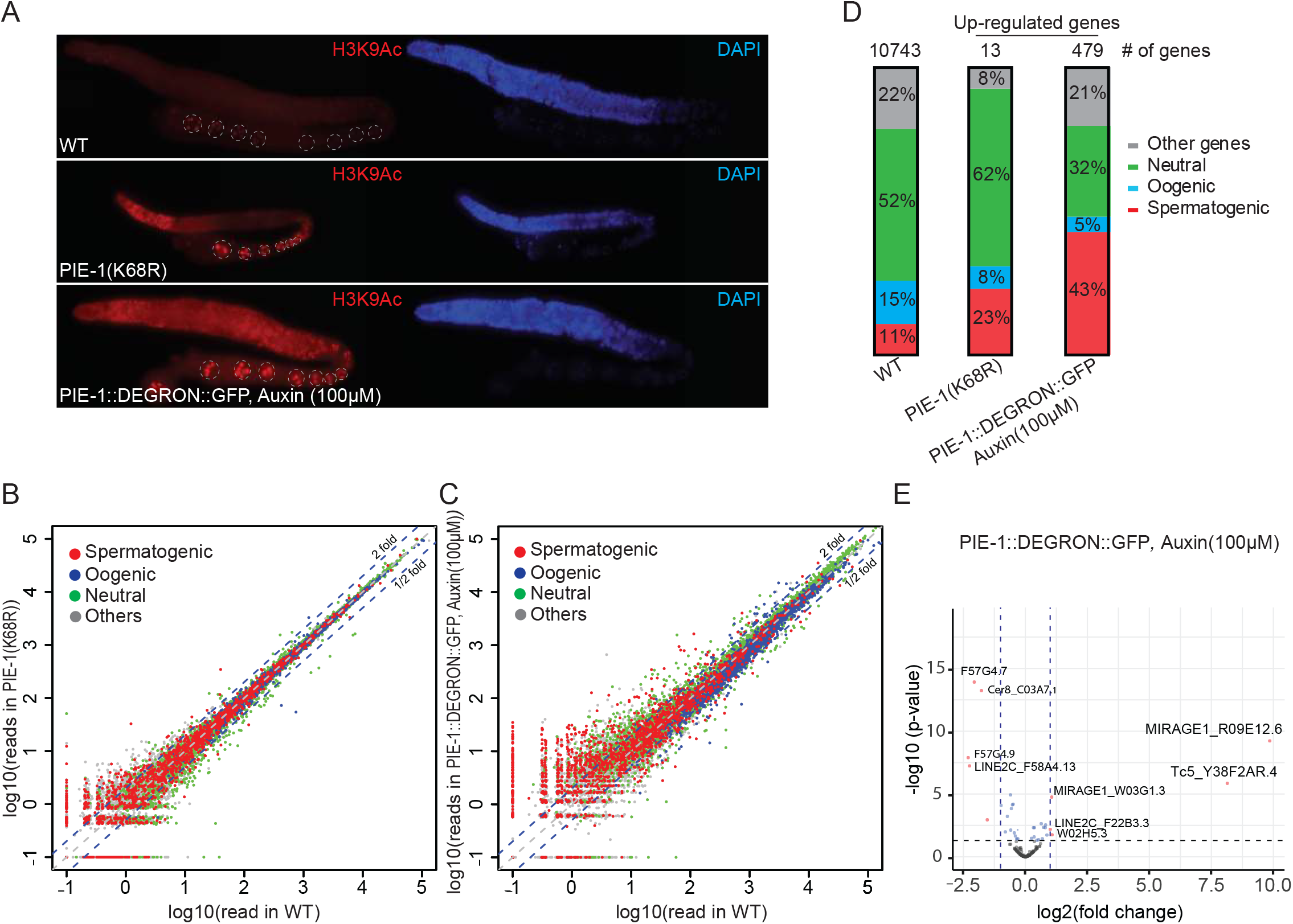
PIE-1 regulates histone H3K9Ac, expression of transposons and spermatogenic genes in the adult germline. (A) Immunofluorescence micrographs of anti-H3K9Ac and DAPI staining in adult gonad of WT, the K68R mutant, and *pie-1::degron* animals (100μM auxin exposure) (B) Scatter plot of reads in *pie-1(ne4303[K68R])* or (C) *pie-1::degron* versus those in WT. Red, blue and green dots are the genes annotated as “Spermatogenic”, “Oogenic”, and “Neutral”, respectively. The green dashed lines indicate two-fold increase or decrease versus WT. A value of 0.1 was assigned to the missing value, thus the ones with a X value of “-1” were not detected in WT. (D) Bar graph shows the composition of each category. “Other genes” are the not-annotated in the previous report (Ortiz et al., 2014). Genes expressed in WT gonad from our mRNA sequencing were used as a reference (“WT gonad”, 10743 genes). The total number of upregulated genes in each mutant is labeled at the top. (E) Volcano plot for the differentially expressed transposons in the *pie-1::degron* mutant. X axis is the fold change versus WT, and Y axis is the adjusted p-value from DESeq2. Dashed lines stand for two-fold change and p-value of 0.05. Transposon’s family name was indicated before its sequence name if available.

Acetylation of H3K9 is associated with active transcription (Peterson and Laniel, 2004). We therefore performed mRNA sequencing on dissected gonads from *pie-1* mutants to determine the extent to which PIE-1 regulates transcription in the adult germline. Whereas *pie-1(ne4303[K68R])* gonads exhibited mild changes in mRNA levels, a group of 479 genes were upregulated by more than 2-fold in *pie-1::degron* depleted gonads, when compared to wild-type (Figures 5B and 5C). Upregulated protein-coding genes included many spermatogenesis-specific genes (Figure 5D), suggesting that PIE-1 promotes the transition from spermatogenesis to oogenesis in the hermaphrodite. Five transposon families were upregulated (Figure 5E), including Tc5 (256-fold) and MIRAGE1 (900-fold), suggesting that PIE-1 activity also promotes transposon silencing in the adult germline.

## Discussion

PIE-1 was first identified as a factor that protects the embryonic germline by preventing somatic differentiation (Mello et al., 1992). We have shown that PIE-1 is also expressed in the adult germline where it engages the Krüppel-type zinc finger protein MEP-1 and the SUMO-conjugating machinery and functions to promote the SUMOylation and activation of the type 1 HDAC, HDA-1 (Figure 6). A parallel study (Kim et al., parallel), extends these findings identifying a role for HDA-1 SUMOylation in promoting association with MEP-1 and the assembly of a NuRD-like chromatin remodeling complex that directs transposon silencing and Argonaute-mediated transcriptional silencing in the germline.

**Figure 6.**
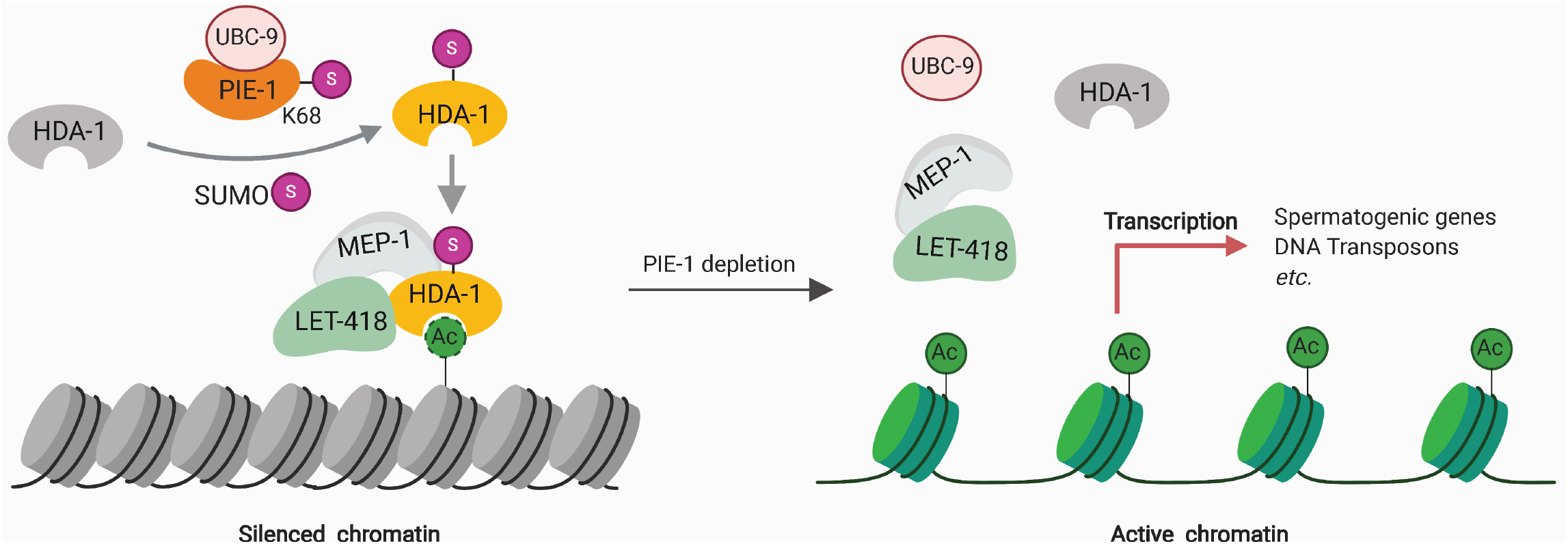
Model. PIE-1 interacts with SUMO pathway components including UBC-9, which leads to PIE-1 SUMOylation. The PIE-1 SUMOylation promotes HDA-1 SUMOylation, which is essential for assembly of MEP-1/LET-418/HDA-1 complex and preserving histone deacetylase activity in the germline.

### MEP-1 assembles functionally distinct complexes regulated by SUMOylation

We found that in the adult germline MEP-1 associates at low levels with HDA-1, and it is this modest association that requires PIE-1 and SUMO pathway factors. These findings suggest that in the adult germline only a small fraction of the HDA-1 protein pool, likely only those molecules that are SUMOylated, can be recruited by MEP-1 for the assembly of a functional NURD complex. Consistent with this idea, a parallel study found that expression of a translational fusion of SUMO (HDA-1::SUMO), from the endogenous *hda-1* locus, rescues an HDA-1 SUMO-acceptor site mutant and dramatically enhances the association of HDA-1`with MEP-1 and other NuRD pathway components (Kim et al., parallel).

Several studies have identified complexes involving MEP-1 or its homologs (Kunert et al., 2009; Passannante et al., 2010; Wu et al., 2012). Studies in *Drosophila* identified a complex, termed MEC, where MEP-1 resides along with Mi-2, but notably, without HDAC (Kunert et al., 2009). In *C. elegans*, MEP-1 was shown to reside in a MEC-like complex in adults, but in embryos this complex was shown to include HDA-1 (a MEC-HDA-1 complex) (Unhavaithaya et al., 2002), but did not appear to contain several other factors normally found in canonical NuRD complexes (Passannante et al., 2010; Wu et al., 2012). In somatic lineages, the remodeling and silencing of germline-specific genes appears to involve a collaboration between the MEC-HDA-1 complex and a conserved DRM (DREAM) complex, comprising DP, Rb, E2F, and MuvB proteins (Harrison et al., 2006). MEC-like and DRM complexes are thought to function together, either transiently or sequentially, on target genes to initiate and maintain transcriptional silencing (Harrison et al., 2006). It’s tempting to speculate that in embryos, perhaps when somatic lineages exit mitosis, changes in Rb phosphorylation promote its interaction with E2F/DP on germline-specific target genes in differentiating somatic cells. The assembly of the DRM complex on these targets may then transiently recruit the MEC-HDA-1 complex to deacetylate and initiate germline chromatin remodeling, and ultimately the recruitment of histone methyltransferases (e.g., PRC2) and methyl-histone binding proteins (e.g., HP1) that maintain silent chromatin.

Interestingly, several components of the SUMO pathway were also identified as factors that prevent germline-gene expression in the *C. elegans* soma (Wu et al., 2012). Suppression of germline gene expression in the soma by MEP-1 and Mi2/LET-418 was shown to be initiated in the embryo (Unhavaithaya et al., 2002). We have shown that HDA-1 is not detectably SUMOylated in the embryo, suggesting that the SUMOylation of other components promotes chromatin remodeling and silencing of germline genes in the embryonic somatic lineages.

In other organisms, several components of the NuRD and DRM complexes have been shown to be SUMOylated or to interact with SUMO. For example, the NuRD complex proteins MTA1 and DCP66 were both identified as SUMO target proteins, and SUMO-interacting motifs in MTA-1 and Mi-2 are thought to promote their functions (Cong et al., 2011; Gong et al., 2006). MEP-1 is SUMOylated both in adults and in embryos (this study), and SUMO-interacting motifs appear to be required for MEP-1 function in the adult germline (Kim et al., parallel). By contrast, SUMOylation of HDA-1 was detected in adults, but not in embryos (Figure 3D). Moreover, mutating the SUMOylation sites of HDA-1 disrupted germline silencing and caused a mortal germline phenotype, but did not disrupt the embryonic functions of MEP-1 (Kim et al., parallel). Thus, SUMOylation of HDA-1 only appears to regulate its functions in the adult germline, suggesting that HDA-1 SUMOylation has cell type, or perhaps cell cycle-specific regulatory effects. Taken together with the finding that other co-factors such as MEP-1 are SUMOylated more broadly, these findings imply that SUMOylation in the context of these chromatin remodeling complexes, does not merely function as a SUMO-glue (Matunis et al., 2006) but rather has specificity depending on which components of the complex are modified and/or when during development.

Genetic studies on vulval induction in *C. elegans* have identified roles for these same factors in controlling the fates of a group of six multipotent hypodermal cells known as vulval precursor cells (VPCs) (Ceol and Horvitz, 2001; Chen and Han, 2001; Horvitz and Sternberg, 1991; Lu and Horvitz, 1998; Solari and Ahringer, 2000; Sulston and Horvitz, 1977; Unhavaithaya et al., 2002; von Zelewsky et al., 2000). The VPCs are born during the L1 stage and arrest division for more than 20 hours until the L3 stage when signaling from the gonadal anchor cell induces the most proximal VPC to adopt a primary vulval cell fate (Sternberg, 2005). The MEC-HDA-1 complex and the DRM co-factors collaborate to ensure that VPCs located farther from the inductive signal stably inactivate their vulval differentiation programs (Ceol and Horvitz, 2001; Chen and Han, 2001; Lu and Horvitz, 1998; Solari and Ahringer, 2000; Unhavaithaya et al., 2002; von Zelewsky et al., 2000). Genetic studies have identified an upstream ETS-domain transcription factor, LIN-1, that functions to repress vulval fates in the VPCs (Beitel et al., 1995; Miley et al., 2004; Tan et al., 1998). In a striking parallel to the findings reported here, LIN-1 was shown to bind UBC-9 and MEP-1 in a yeast 2-hybrid screen, and mutations in LIN-1 consensus SUMO acceptor lysine residues caused partial loss-of-function phenotypes and reduced association with MEP-1 (Leight et al., 2005). SUMOylation of LIN-1 therefore appears to promote its activity and to stimulate recruitment of the MEP-1/HDA-1/MEC-like complex to mediate transcriptional silencing of LIN-1 targets in the VPCs (Leight et al., 2005). It is intriguing that the VPCs undergo this regulation during an extended G1 arrest, raising the possibility that LIN-1 and PIE-1 provide analogous functions as SUMO E3-like or substrate-bridging factors, needed to overcome a cell cycle-dependent regulation that inhibits the association of HDA-1 with MEP-1 and other NuRD pathway components. It will be interesting to explore whether HDA-1 is a target of LIN-1 dependent SUMOylation or if other targets mediate this regulation.

### SUMOylation promotes stable transitions in gene expression and cellular differentiation

We have shown that the SUMO pathway promotes PIE-1 activity in both the adult and embryonic germline. In embryos, our genetic studies implicate SUMO in the previously described role for PIE-1 in protecting embryonic germline blastomeres from adopting somatic cell fates. This PIE-1 function is thought to involve the suppression of pol-II activity in the germline (Seydoux and Dunn, 1997), perhaps through inhibition of the CTD kinase (Batchelder et al., 1999; Ghosh and Seydoux, 2008). How SUMO promotes this inhibition will require further study, for as we have shown, PIE-1 itself is not detectably SUMOylated in early embryos. This apparent paradox could be explained by at least two scenarios that are not mutually exclusive. PIE-1 and SUMO might act together in the adult germline—prior to fertilization—to, for example, prepare the embryonic chromatin for maintenance of germline fates in the embryo. Alternatively, PIE-1 and SUMO could act separately in the embryonic germline, where SUMO may regulate targets independently of PIE-1 SUMOylation. Perhaps more consistent with this latter possibility, we noted a change in the genetic relationship between PIE-1 and the SUMO pathway during embryogenesis. We found that the SUMO E3 ligase GEI-17—the Su(var)2-10 homolog— was dispensable for PIE-1- and SUMO-dependent silencing in the adult germline (Kim et al., parallel), but was required along with PIE-1 activity for proper specification of the embryonic germline blastomere P2. Depletion of *gei-17* in *pie-1* heterozygous worms caused nearly 100% of embryos to die without germ cells and with extra intestine, similar to the phenotype of dead embryos made by *pie-1* homozygous worms. Thus PIE-1 collaborates with GEI-17 and the SUMO pathway to protect the embryonic germ cell fate. One picture that emerges from this work is that SUMO plays diverse roles in regulating protein interactions and activities during development, and frequently appears to promote stable transitions to new patterns of gene expression and cellular differentiation. For example, in the adult hermaphrodite germline, we have shown that SUMO and PIE-1 function together to prevent the expression of spermatogenesis genes after the transition to oogenesis. In embryos they function together to prevent embryonic somatic cells from reverting to patterns of germline-specific gene expression, and as discussed above to prevent germline blastomeres from adopting somatic cell fates. The SUMO proteomics described here, along with the remarkable ease of CRISPR genome editing in *C. elegans*, should enable a very detailed look at specific SUMOylation events and how they regulate differentiation and other cellular mechanisms.

## Materials and Methods

### C. elegans strains and genetics

Strains and alleles used in this study were listed in Table S3. Worms were, unless otherwise stated, cultured at 20°C on NGM plates seeded with OP50 *Escherichia coli*, and genetic analyses were performed essentially as described (Brenner, 1974).

### CRISPR/Cas9 genome editing

The Co-CRISPR strategy (Kim et al., 2014) using *unc-22* sgRNA as a co-CRISPR marker was used to enrich HR events for tagging a gene of interest with the non-visualizing epitope (*6xhis* and *10xhis*) or introduction of a point mutation (G56R). To screen for insertions of *6xhis* and *10xhis*, we used two-round PCR: the First PCR was performed with primers (F: cctcaaaaaccaagcgaaaacc R: ccggctgctatttcattgat), and 1 μl of the 1^st^ PCR product was used as a template for the 2^nd^ PCR with primers (F:gagactcccgctataaacga R:ctcaagcaggcgacaacgcc). To detect *6x*/*10xhis* knock-ins, the final products were run either on 2% Tris/borate/EDTA (TBE) gel or 10% PAGE gel. *The ubc-9(G56R)* mutation introduced an *Hae*III restriction fragment length polymorphisms (RFLP) that was used to screen for G56R conversion in PCR products (F: cattacatggcaagtcggg, R: cgttgccgcatacagaaatag). For visualization of either GFP tag, F1 rollers were mounted under coverslips on 2% agarose pads to directly screen for GFP expressing animals as described previously (Kim et al., 2014). *3xflag* knock-ins to *pie-1(ne4303)* were screened by PCR using previously reported primers (Kim et al., 2014). sgRNA construct: previously generated *pie-1* sgRNA plasmid (Kim et al., 2014) was used for *pie-1::degron::gfp* and *pie-1(K68R)::*flag. Others were constructed by ligating annealed sgRNA oligonucleotides to *Bsa*l-cut pRB1017 (Arribere et al., 2014) and sgRNA sequences were listed in Table S4.

Donor template: unless otherwise stated, a silent mutation to disrupt the PAM site in each HR donor was introduced by PCR sewing.

*ubc-9(G56R)*: for the *ubc-9(G56R)* donor construct, a *ubc-9* fragment was amplified with primers (F: cattacatggcaagtcggg, R: gacgactaccacgaagcaagc) and this fragment was cloned into the Blunt II-TOPO vector (Thermo Fisher Scientific, K2800-20). To introduce the point mutation (G56R) and mutate the seeding region, overlap extension PCR was used. *6xhis::smo-1*/*10xhis::smo-1*: using PCR sewing, either *6xhis* (caccatcaccaccatcac) or *10xhis* fragment (caccatcaccatcaccatcaccaccatcac) was introduced immediately after the start codon in the previously generated *smo-1* fragment (Kim et al., 2014). The resulting PCR product was cloned into the Blunt II-TOPO vector. Tagging with *his* tag on the N-terminus of *smo-1* disrupted the PAM site.

*mep-1::gfp::tev::3xflag*: for the *mep-1::gfp::tev::3xflag* donor construct, a *mep-1* fragment was amplified with primers (F1:gaaattcgctggcagtttct R1: ctgcaacttcgatcaatcga) from N2 genomic DNA and inserted into the pCR-Blunt II-TOPO vector. Overlap extension PCR was used to introduce a *Nhe*I site immediately before the stop codon in this *mep-1* fragment. The *Nhe*I site was used to insert the *gfp::tev::3xflag* coding sequence.

*pie-1(K68R)::3xflag* and *pie-1::degron::gfp*: previously generated donor plasmid (Kim et al., 2014) was used for *pie-1(K68R)::3xflag*. To introduce the degron sequence (Zhang et al., 2015) into *pie-1::gfp* donor plasmid previously built in pCR-Blunt II TOPO (Kim et al., 2014), PCR was performed with primers including *NheI* site using pLZ29 plasmid (Zhang et al., 2015). The PCR products were cut with *NheI* restriction enzyme, gel-purified and cloned into the pCR-Blunt II Topo vector.

### Yeast two-hybrid analysis

The yeast two-hybrid screen was performed by Hybrigenics services (Paris, France, http://www/hybrigenics-services.com). The coding sequence for amino acids 2–335 of *C. elegans pie-1* (NM_001268237.1) was amplified by PCR from N2 cDNA and cloned into pB66 via C-terminal fusion with the Gal4 DNA-binding domain (Gal-4-PIE-1). The construct was checked by sequencing and used as a bait to screen a random-primed *C. elegans* Mixed Stages cDNA library constructed into pP6. 5 million clones (5-fold the complexity of the library). The library was screened using a mating approach with YHGX13 (Y187 ade2-101::loxp-kanMX-loxP, matα) and CG1945 (mata) yeast strains as previously described (Fromont-Racine et al., 1997). 153 His+ colonies were selected on a medium lacking tryptophan, leucine, and histidine, and supplemented with 0.5 mM 3-aminotriazole to prevent bait autoactivation. The prey fragments of the positive clones were amplified by PCR and sequenced at their 5’ and 3’ junctions. The resulting sequences were used to identify the corresponding interacting proteins in the GenBank database (NCBI) using a fully automated procedure. A confidence score (predicted biological score [PBS]) was attributed to each interaction as previously described (Formstecher et al., 2005).

### Auxin Treatment

The *pie-1::degron::gfp* strain was introduced into CA1199 (*unc-119(ed3); ieSi38 [Psun-1::TIR1::mRuby::sun-1 3’UTR, cb-unc-119(+)] IV)* (Zhang et al., 2015) by CRISPR/Cas9-mediated genome editing. The degron tagged-L1 larval stage worms were plated on NGM plate containing 100 μM auxin indole-3-acetic acid (IAA) (Alfa Aesar, A10556). Worms were collected for gonad dissection when the L1 reached adult stages.

### RNAi

RNAi was performed by feeding worms *E. coli* strain HT115 (DE3) transformed with the control vector or a gene-targeting construct from the *C. elegans* RNAi Collection (Kamath and Ahringer, 2003). For the genetic analysis, L4 larval stage animals were placed on RNAi plates (NGM plates containing 1mM isopropyl β-d-thiogalactoside (IPTG) and 100 μg/ml ampicillin seeded with dsRNA-containing bacteria) and allowed to develop into adults. After 24h, adult animals were transferred to fresh RNAi plates and allowed to lay eggs overnight. On the following day, the unhatched eggs were analyzed 12 h later after adults were removed from the test plates. For the biochemical assay (large culture), ~200,000 synchronous L4 worms were placed on RNAi plates including 0.4mM IPTG, 100 μg/ml ampicillin, and 12.5 μg/ml tetracycline.

### Immunofluorescence

Gonads were dissected on glass slide (Thermo Fisher Scientific, 1256820) in Egg Buffer (25mM HEPES pH 7.5, 118mM NaCl, 48mM KCl, 2 mM CaCl_2_, 2mM MgCl_2_) containing 0.2mM tetramisole (Sigma-Aldrich, L9756). The samples were transfer into slickseal^TM^ microcentrifuge tube (National Scientific, CA170S-BP) and fixed with 2% paraformaldehyde (Electron Microscopy Science, Nm15710) in phosphate-buffered saline (PBS) pH 7.4 for 10min, and −20°C cold 100% methanol for 5min. After fixation, samples were washed four times with PBST (PBS containing 0.1% Tween20) containing 0.1% bovine serum albumin (BSA), blocked in PBST containing 1% BSA for 1hr, and then incubated with primary antibody in PBST containing 1% BSA overnight at 4°C. After four washes with PBST containing 0.1% BSA, secondary antibodies were applied in PBST containing 1% BSA for 2h at room temperature. Sample were washed four times with PBST containing 0.1% BSA, transferred to Poly-L-lysine coated slides (Labscientific, 7799) and mounted with 10 μl of SlowFade Diamond Antifade Mountant with DAPI (Life Technologies, S36964). The primary antibodies (1:100) used was anti-acetyl-histone H3Lys9 (Abcam, ab12179). The secondary antibodies (1:1000) used were: goat anti-mouse IgG(H+L) Alexa Fluor 594 (Thermo Fisher Scientific, A-11005).

### Microscopy

For live imaging, worms and embryos were mounted in M9 on a 2% agarose pad with or without 1mM tetramisole. Epi-fluorescence and differential interference contrast (DIC) microscopy were performed using an Axio Imager M2 Microscope (Zeiss). Images were captured with an ORCA-ER digital camera (Hamamatsu) and processed using Axiovision software (Zeiss). Confocal images were acquired using a Zeiss Axiover 200M microscope equipped with a Yokogawa CSU21 spinning disk confocal scan head and custom laser launch and relay optics (Solamere Technology Group). Stacks of images were taken and analyzed using ImageJ.

### Immunoprecipitation

Either synchronous adult worms (~200,000 animals) or early embryos (bleached from 10×100,000 adult animals) were collected and washed three times with M9 buffer before being homogenized in lysis buffer [20mM HEPES pH 7.5, 125mM Na_3_C_6_H_5_O_7_ (sodium citrate), 0.1%(v/v) Tween 20, 0.5%(v/v) Triton X-100, 2mM MgCl_2_, 1mM DTT, and a Mini Protease Inhibitor Cocktail Tablet (Roche)] using a FastPrep-24 benchtop homogenizer (MP Biomedicals). Worm or embryo lysates were centrifuged twice at 14,000×g for 30 min at 4°C, and supernatants were incubated with GFP-binding protein (GBP) beads for 1.5 h at 4°C on a rotating shaker. The beads were washed three times with immunoprecipitation (IP) buffer containing protease inhibitor for 5 min each wash and then washed twice with high-salt wash buffer (50mM HEPES pH 7.5, 500mM KCl, 0.05% NP40, 0.5mM DTT, and protease inhibitor). Immune complexes were eluted with elution buffer (50mM Tris-Cl pH 8.0, 1xSDS) for 10 min at 95°C.

### Affinity chromatography of histidine-tagged SUMO

Synchronous adult worms (~200,000) or 500 μl of packed embryos (bleached from (~1,000,000 synchronous adult worms) were homogenized in lysis buffer at pH 8.0 (6M guanidine-HCl, 100mM Na_2_HPO_4_/NaH_2_PO_4_ pH 8.0, and 10mM Tris-HCl pH8.0) using a FastPrep-24 benchtop homogenizer (MP Biomedicals). Lysates were cleared by centrifugation at 14,000×g for 30 min at 4°C and equalized using quick start Bradford 1x dye reagent (BioRad, 5000205). Ni-NTA resin was washed three times with lysis buffer containing 20mM imidazole pH 8.0 and 5mM β-mercaptoethanol while samples were prepared. To equalized samples, we added imidazole pH 8.0 to 20mM and β-mercaptoethanol to 5mM, and then the samples were incubated with 100 μl of pre-cleared 50% slurry of Ni-NTA resin (Qiagen, 30210) for 2–3 h at room temperature on a rotating shaker. Ni-NTA resin was washed in 1-ml aliquots of the following series of buffers: Buffer 1 pH 8.0 (6M guanidine-HCl, 100mM Na_2_HPO_4_/NaH_2_PO_4_ pH 8.0, 10mM Tris-HCl pH 8.0, 10mM imidazole pH 8.0, 5mM β-mercaptoethanol, and 0.1% Triton X-100), Buffer 2 pH 8.0 (8M urea, 100mM Na_2_HPO_4_/NaH_2_PO_4_ pH 8.0, 10mM Tris-HCl pH 8.0, 10mM imidazole pH 8.0, 5mM β-mercaptoethanol, and 0.1% Triton X-100), Buffer 3 pH 7.0 (8M urea, 100mM Na_2_HPO_4_/NaH_2_PO_4_ pH 7.0, 10mM Tris-HCl pH 7.0, 10mM imidazole pH 7.0, 5mM β-mercaptoethanol, and 0.1% Triton X-100), Buffer 4 pH 6.3 (8M urea, 100mM Na_2_HPO_4_/NaH_2_PO_4_ pH6.3, 10mM Tris-HCl pH 6.3, 10mM imidazole pH 6.3, 5mM β-mercaptoethanol, and 0.1% Triton X-100), Buffer 5 pH 6.3 (8M urea, 100mM Na_2_HPO_4_/NaH_2_PO_4_ pH 6.3, 10mM Tris-HCl pH 6.3, and 5mM β-mercaptoethanol). To purify the polyhistidine-tagged proteins under denaturing conditions, 8M urea was added to all wash buffers. Triton-X-100, the non-ionic detergent was used to reduce nonspecific hydrophobic interactions. Imidazole (10mM) was used to increase the stringency of the wash by reducing nonspecific protein binding to the resin. The use of wash buffers with gradually decreasing pH (pH8 to pH6.3) also reduced nonspecific binding of proteins by protonating the neutral histidine and thereby removing the weakly bound proteins that may contain tandem repeats of the histidine. The SUMOylated proteins were eluted with elution buffer pH 7.0 (7M urea, 100mM Na_2_HPO_4_/NaH_2_PO_4_ pH 7.0, 10mM Tris-HCl pH 7.0, and 500mM imidazole pH 7.0) For western blotting, input samples containing guanidine-HCl were diluted with H_2_O (1:6) and then purified by trichloroacetic acid (TCA) precipitation: an equal volume of 10% TCA was added to diluted samples, which were then incubated on ice for 20 min and centrifuged for 20 min at 4°C; the obtained pellet was washed with 100 μl of ice-cold ethanol and then resuspended in Tris-HCl buffer pH 8.0.

### Western blot analysis

NuPAGE LDS sample buffer (4x) (Thermo Fisher Scientific, NP0008) was added to samples, which were then loaded on precast NuPAGE Novex 4-12% Bis-Tris protein gel (Life Technologies, NP0321BOX) and transferred onto a polyvinylidene difluoride (PVDF) membrane (Bio-Rad Laboratories, 1704157) using Mini Trans-Blot cells (Bio-Rad Laboratories, 1703930) at 80 V for 2.2h at 4°C. Membranes were blocked with 5% skim milk and probed with primary antibodies: anti-FLAG (1:1000) (Sigma-Aldrich, F1804), anti-MRG-1 (1:1000) (Novus Biologicals, 49130002), anti-HDA-1 (1:2500) (Novus Biologicals, 38660002), anti-LET-418 (1:1000) (Novus Biologicals, 48960002), anti-SMO-1 (1:1000) (purified from Hybridoma cell cultures, the Hybridoma cell line was gift from Ronald T. Hay, University of Dundee)(Pelisch et al., 2014), anti-tubulin (1:2000) (Bio-Rad, MCA77G), anti-PIE-1(P4G5) (1:100), and anti-PRG-1(1:1000). Antibody binding was detected with secondary antibodies: goat anti-mouse (1:2500) (Thermo Fisher Scientific, 62-6520), mouse anti-rabbit (1:3000) (Abcam, ab99697), anti-rat (1:5000) (Jackson ImmunoResearch Labs, 712-035-150).

### RNA-seq

Germline RNA was extracted from 100 dissected gonads using TRI reagent (Sigma Aldrich, T9424). Total RNA (500ng per replica) was used for library production using KAPA RNA HyperPrep with RiboErase (Kapa Biosystems, KK8560) and KAPA single-indexed adapter (Kapa Biosystems, KK8700) for Illumina platforms. The RNA libraries were sequenced at pair-end on NextSeq 500 Sequencer with Illumina NextSeq 500/550 high output kit v2.5 (150 cycles) (Illumina, 20024907). Salmon was used to map and quantify the reads against the worm database WS268, and its output files were imported to DESeq2 in R. The differentially expressed genes were defined as two-fold change and adjusted p-value less than 0.05 (Table S5). The scatter plots were generated by the plot function in R. Comparisons between repeats of each sample are in Figure S5.

### Mass Spectrometry Sample Preparation

The precipitated His-SMO-1 pull down samples are resolubilized in 8 M Urea, 100 mM Tris, pH 8.0 followed by reduction with 5 mM TCEP for 20 min, alkylation with 10 mM iodoacetamide for 15 min and trypsin digestion. Each fraction was analyzed on a Q Exactive HF mass spectrometer (Thermo Fisher Scientific) coupled to a nano UHPLC Easy-nLC 1000 via a nano-electrospray ion source. Peptides were separated on a home-packed capillary reverse phase column (75-μm internal diameter × 15 cm of 1.8 μm, 120 Å UHPLC-XB-C18 resin) with a 110-min gradient of A and B buffers (buffer A, 0.1% formic acid; buffer B, 100% ACN/0.1% formic acid). A lock mass of 445.120025 m/z was used for internal calibration. Electrospray ionization was carried out at 2.0 kV, with the heated capillary temperature set to 275 °C. Full-scan mass spectra were obtained in the positive-ion mode over the m/z range of 300 to 2000 at a resolution of 120,000. MS/MS spectra were acquired in the Orbitrap for the 15 most abundant multiply-charged species in the full-scan spectrum having signal intensities of > 1 × 10^−5^ NL at a resolution of 15,000. Dynamic exclusion was set such that MS/MS was acquired only once for each species over a period of 30 sec. For His-SMO-1 pull down protein identification, the MS data was searched against the concatenated forward and reversed *C. elegans* protein database (WS233) using by Prolucid (Xu et al., 2006) and DTASelect 2 (Tabb et al., 2002) (≤ 1% FDR at the peptide level).

## Acknowledgement

We thank members of Mello and Ambros labs for discussions. C.C.M. is a Howard Hughes Medical Institute Investigator.

## Author contributions

Conceptualization, H.K., D.C., and C. C. M.; Investigation, H.K., Y.D., S.L., M.Z., M.D., and C.C.M.; Writing-Original draft, H.K., and C.C.M.; Writing-Review & Editing, H.K., Y.D., D.C., and C. C. M.; Supervision, C.C.M. and M.D.

## References

Arribere, J.A., Bell, R.T., Fu, B.X., Artiles, K.L., Hartman, P.S., and Fire, A.Z. (2014). Efficient marker-free recovery of custom genetic modifications with CRISPR/Cas9 in Caenorhabditis elegans. Genetics 198, 837–846.

Batchelder, C., Dunn, M.A., Choy, B., Suh, Y., Cassie, C., Shim, E.Y., Shin, T.H., Mello, C., Seydoux, G., and Blackwell, T.K. (1999). Transcriptional repression by the Caenorhabditis elegans germ-line protein PIE-1. Genes Dev 13, 202–212.

Beitel, G.J., Tuck, S., Greenwald, I., and Horvitz, H.R. (1995). The Caenorhabditis elegans gene lin-1 encodes an ETS-domain protein and defines a branch of the vulval induction pathway. Genes Dev 9, 3149–3162.

Betting, J., and Seufert, W. (1996). A yeast Ubc9 mutant protein with temperature-sensitive in vivo function is subject to conditional proteolysis by a ubiquitin- and proteasome-dependent pathway. J Biol Chem 271, 25790–25796.

Beurton, F., Stempor, P., Caron, M., Appert, A., Dong, Y., Chen, R.A., Cluet, D., Coute, Y., Herbette, M., Huang, N., et al. (2019). Physical and functional interaction between SET1/COMPASS complex component CFP-1 and a Sin3S HDAC complex in C. elegans. Nucleic Acids Res 47, 11164–11180.

Blackshear P.J., P.R.S., Lai W.S. (2005). Tandem CCCH Zinc Finger Proteins in mRNA Binding. In Iuchi S, Kuldell N (eds) Zinc Finger Proteins: from atomic contact to cellular function Molecular biology intelligence unit (Springer, Boston, MA).

Brenner, S. (1974). The genetics of Caenorhabditis elegans. Genetics 77, 71–94.

Broday, L., Kolotuev, I., Didier, C., Bhoumik, A., Gupta, B.P., Sternberg, P.W., Podbilewicz, B., and Ronai, Z. (2004). The small ubiquitin-like modifier (SUMO) is required for gonadal and uterine-vulval morphogenesis in Caenorhabditis elegans. Genes Dev 18, 2380–2391.

Capili, A.D., and Lima, C.D. (2007). Taking it step by step: mechanistic insights from structural studies of ubiquitin/ubiquitin-like protein modification pathways. Curr Opin Struct Biol 17, 726–735.

Ceol, C.J., and Horvitz, H.R. (2001). dpl-1 DP and efl-1 E2F act with lin-35 Rb to antagonize Ras signaling in C. elegans vulval development. Mol Cell 7, 461–473.

Chen, Z., and Han, M. (2001). C. elegans Rb, NuRD, and Ras regulate lin-39-mediated cell fusion during vulval fate specification. Curr Biol 11, 1874–1879.

Cong, L., Pakala, S.B., Ohshiro, K., Li, D.Q., and Kumar, R. (2011). SUMOylation and SUMO-interacting motif (SIM) of metastasis tumor antigen 1 (MTA1) synergistically regulate its transcriptional repressor function. J Biol Chem 286, 43793–43808.

Formstecher, E., Aresta, S., Collura, V., Hamburger, A., Meil, A., Trehin, A., Reverdy, C., Betin, V., Maire, S., Brun, C., et al. (2005). Protein interaction mapping: a Drosophila case study. Genome Res 15, 376–384.

Fromont-Racine, M., Rain, J.C., and Legrain, P. (1997). Toward a functional analysis of the yeast genome through exhaustive two-hybrid screens. Nat Genet 16, 277–282.

Geiss-Friedlander, R., and Melchior, F. (2007). Concepts in sumoylation: a decade on. Nat Rev Mol Cell Biol 8, 947–956.

Ghosh, D., and Seydoux, G. (2008). Inhibition of transcription by the Caenorhabditis elegans germline protein PIE-1: genetic evidence for distinct mechanisms targeting initiation and elongation. Genetics 178, 235–243.

Gill, G. (2004). SUMO and ubiquitin in the nucleus: different functions, similar mechanisms? Genes Dev 18, 2046–2059.

Gong, Z., Brackertz, M., and Renkawitz, R. (2006). SUMO modification enhances p66-mediated transcriptional repression of the Mi-2/NuRD complex. Mol Cell Biol 26, 4519–4528.

Harrison, M.M., Ceol, C.J., Lu, X., and Horvitz, H.R. (2006). Some C. elegans class B synthetic multivulva proteins encode a conserved LIN-35 Rb-containing complex distinct from a NuRD-like complex. Proc Natl Acad Sci U S A 103, 16782–16787.

Hendriks, I.A., and Vertegaal, A.C. (2016). A comprehensive compilation of SUMO proteomics. Nat Rev Mol Cell Biol 17, 581–595.

Holway, A.H., Hung, C., and Michael, W.M. (2005). Systematic, RNA-interference-mediated identification of mus-101 modifier genes in Caenorhabditis elegans. Genetics 169, 1451–1460.

Horvitz, H.R., and Sternberg, P.W. (1991). Multiple intercellular signalling systems control the development of the Caenorhabditis elegans vulva. Nature 351, 535–541.

Johnson, E.S. (2004). Protein modification by SUMO. Annu Rev Biochem 73, 355–382.

Jones, D., Crowe, E., Stevens, T.A., and Candido, E.P. (2002). Functional and phylogenetic analysis of the ubiquitylation system in Caenorhabditis elegans: ubiquitin-conjugating enzymes, ubiquitin-activating enzymes, and ubiquitin-like proteins. Genome Biol 3, RESEARCH0002.

Kamath, R.S., and Ahringer, J. (2003). Genome-wide RNAi screening in Caenorhabditis elegans. Methods 30, 313–321.

Kim, H., Ishidate, T., Ghanta, K.S., Seth, M., Conte, D., Jr., Shirayama, M., and Mello, C.C. (2014). A co-CRISPR strategy for efficient genome editing in Caenorhabditis elegans. Genetics 197, 1069–1080.

Kunert, N., Wagner, E., Murawska, M., Klinker, H., Kremmer, E., and Brehm, A. (2009). dMec: a novel Mi-2 chromatin remodelling complex involved in transcriptional repression. EMBO J 28, 533–544.

Leight, E.R., Glossip, D., and Kornfeld, K. (2005). Sumoylation of LIN-1 promotes transcriptional repression and inhibition of vulval cell fates. Development 132, 1047–1056.

Lim, Y., Lee, D., Kalichamy, K., Hong, S.E., Michalak, M., Ahnn, J., Kim, D.H., and Lee, S.K. (2014). Sumoylation regulates ER stress response by modulating calreticulin gene expression in XBP-1-dependent mode in Caenorhabditis elegans. Int J Biochem Cell Biol 53, 399–408.

Lu, X., and Horvitz, H.R. (1998). lin-35 and lin-53, two genes that antagonize a C. elegans Ras pathway, encode proteins similar to Rb and its binding protein RbAp48. Cell 95, 981–991.

Matunis, M.J., Zhang, X.D., and Ellis, N.A. (2006). SUMO: the glue that binds. Dev Cell 11, 596–597.

Mello, C.C., Draper, B.W., Krause, M., Weintraub, H., and Priess, J.R. (1992). The pie-1 and mex-1 genes and maternal control of blastomere identity in early C. elegans embryos. Cell 70, 163–176.

Mello, C.C., Schubert, C., Draper, B., Zhang, W., Lobel, R., and Priess, J.R. (1996). The PIE-1 protein and germline specification in C. elegans embryos. Nature 382, 710–712.

Miley, G.R., Fantz, D., Glossip, D., Lu, X., Saito, R.M., Palmer, R.E., Inoue, T., Van Den Heuvel, S., Sternberg, P.W., and Kornfeld, K. (2004). Identification of residues of the Caenorhabditis elegans LIN-1 ETS domain that are necessary for DNA binding and regulation of vulval cell fates. Genetics 167, 1697–1709.

Ortiz, M.A., Noble, D., Sorokin, E.P., and Kimble, J. (2014). A new dataset of spermatogenic vs. oogenic transcriptomes in the nematode Caenorhabditis elegans. G3 (Bethesda) 4, 1765–1772.

Passannante, M., Marti, C.O., Pfefferli, C., Moroni, P.S., Kaeser-Pebernard, S., Puoti, A., Hunziker, P., Wicky, C., and Muller, F. (2010). Different Mi-2 complexes for various developmental functions in Caenorhabditis elegans. PLoS One 5, e13681.

Pelisch, F., Sonneville, R., Pourkarimi, E., Agostinho, A., Blow, J.J., Gartner, A., and Hay, R.T. (2014). Dynamic SUMO modification regulates mitotic chromosome assembly and cell cycle progression in Caenorhabditis elegans. Nat Commun 5, 5485.

Peterson, C.L., and Laniel, M.A. (2004). Histones and histone modifications. Curr Biol 14, R546–551.

Prendergast, J.A., Ptak, C., Arnason, T.G., and Ellison, M.J. (1995). Increased ubiquitin expression suppresses the cell cycle defect associated with the yeast ubiquitin conjugating enzyme, CDC34 (UBC3). Evidence for a noncovalent interaction between CDC34 and ubiquitin. J Biol Chem 270, 9347–9352.

Psakhye, I., and Jentsch, S. (2012). Protein group modification and synergy in the SUMO pathway as exemplified in DNA repair. Cell 151, 807–820.

Reese, K.J., Dunn, M.A., Waddle, J.A., and Seydoux, G. (2000). Asymmetric segregation of PIE-1 in C. elegans is mediated by two complementary mechanisms that act through separate PIE-1 protein domains. Mol Cell 6, 445–455.

Reichman, R., Shi, Z., Malone, R., and Smolikove, S. (2018). Mitotic and Meiotic Functions for the SUMOylation Pathway in the Caenorhabditis elegans Germline. Genetics 208, 1421–1441.

Rothbauer, U., Zolghadr, K., Muyldermans, S., Schepers, A., Cardoso, M.C., and Leonhardt, H. (2008). A versatile nanotrap for biochemical and functional studies with fluorescent fusion proteins. Mol Cell Proteomics 7, 282–289.

Santella, A., Kovacevic, I., Herndon, L.A., Hall, D.H., Du, Z., and Bao, Z. (2016). Digital development: a database of cell lineage differentiation in C. elegans with lineage phenotypes, cell-specific gene functions and a multiscale model. Nucleic Acids Res 44, D781–785.

Seydoux, G., and Braun, R.E. (2006). Pathway to totipotency: lessons from germ cells. Cell 127, 891–904.

Seydoux, G., and Dunn, M.A. (1997). Transcriptionally repressed germ cells lack a subpopulation of phosphorylated RNA polymerase II in early embryos of Caenorhabditis elegans and Drosophila melanogaster. Development 124, 2191–2201.

Shen, T.H., Lin, H.K., Scaglioni, P.P., Yung, T.M., and Pandolfi, P.P. (2006). The mechanisms of PML-nuclear body formation. Mol Cell 24, 331–339.

Shi, Y., and Mello, C. (1998). A CBP/p300 homolog specifies multiple differentiation pathways in Caenorhabditis elegans. Genes Dev 12, 943–955.

Solari, F., and Ahringer, J. (2000). NURD-complex genes antagonise Ras-induced vulval development in Caenorhabditis elegans. Curr Biol 10, 223–226.

Sternberg, P.W. (2005). Vulval development. WormBook, 1–28.

Strome, S., and Updike, D. (2015). Specifying and protecting germ cell fate. Nat Rev Mol Cell Biol 16, 406–416.

Sulston, J.E., and Horvitz, H.R. (1977). Post-embryonic cell lineages of the nematode, Caenorhabditis elegans. Dev Biol 56, 110–156.

Tabb, D.L., McDonald, W.H., and Yates, J.R., 3rd (2002). DTASelect and Contrast: tools for assembling and comparing protein identifications from shotgun proteomics. J Proteome Res 1, 21–26.

Tan, P.B., Lackner, M.R., and Kim, S.K. (1998). MAP kinase signaling specificity mediated by the LIN-1 Ets/LIN-31 WH transcription factor complex during C. elegans vulval induction. Cell 93, 569–580.

Tatham, M.H., Rodriguez, M.S., Xirodimas, D.P., and Hay, R.T. (2009). Detection of protein SUMOylation in vivo. Nat Protoc 4, 1363–1371.

Tenenhaus, C., Schubert, C., and Seydoux, G. (1998). Genetic requirements for PIE-1 localization and inhibition of gene expression in the embryonic germ lineage of Caenorhabditis elegans. Dev Biol 200, 212–224.

Tenenhaus, C., Subramaniam, K., Dunn, M.A., and Seydoux, G. (2001). PIE-1 is a bifunctional protein that regulates maternal and zygotic gene expression in the embryonic germ line of Caenorhabditis elegans. Genes Dev 15, 1031–1040.

Unhavaithaya, Y., Shin, T.H., Miliaras, N., Lee, J., Oyama, T., and Mello, C.C. (2002). MEP-1 and a homolog of the NURD complex component Mi-2 act together to maintain germline-soma distinctions in C. elegans. Cell 111, 991–1002.

von Zelewsky, T., Palladino, F., Brunschwig, K., Tobler, H., Hajnal, A., and Muller, F. (2000). The C. elegans Mi-2 chromatin-remodelling proteins function in vulval cell fate determination. Development 127, 5277–5284.

Wu, X., Shi, Z., Cui, M., Han, M., and Ruvkun, G. (2012). Repression of germline RNAi pathways in somatic cells by retinoblastoma pathway chromatin complexes. PLoS Genet 8, e1002542.

Xu, T., Venable, J.D., Park, S.K., Cociorva, D., Lu, B., Liao, L., Wohlschlegel, J., Hewel, J., and Yates, J.R., 3rd (2006). ProLuCID, a fast and sensitive tandem mass spectra-based protein identification program. Mol Cell Proteomics 5(10), S174.

Zhang, L., Ward, J.D., Cheng, Z., and Dernburg, A.F. (2015). The auxin-inducible degradation (AID) system enables versatile conditional protein depletion in C. elegans. Development 142, 4374–4384.

