## Supplemental figures for "PIE-1 promotes SUMOylation and activation of HDAC1 during the *C. elegans* oogenesis"

Figure S1 PIE-1 is insoluble and unstable

(A) PIE-1 solubility in 9 different lysis buffer (B1-B9) showed PIE-1 is insoluble, compared to other proteins, PRG-1 and tubulin. S and P represent supernatant and Pellet, respectively.

(B) Silver staining of purified PIE-1::GFP revealed a cleavage fragment.

(C) MS analysis identified S327 is the cleavage site for PIE-1.

Figure S2 SUMO purification using Ni-NTA column in *his* tagged *smo-1* animal

(A) and (B) western blot comparing purified SUMO target proteins between with his6::smo-1 in (A) and with his10::smo-1 in (B).

(C) Silver staining gel and western blot showing procedure of SUMO purification and purified SUMO proteins.

Figure S3 PIE-1 expression in the adult germline and during embryogenesis

Confocal images of adult germline and embryos in a strain expressing PIE-1::GFP.

Figure S4 Auxin-induced depletion of PIE-1::DEGRON::GFP

Live images of adult animals expressing PIE-1::DEGRON::GFP and *sun-1* promoter-driven TIR1 without auxin in (A) with auxin in (B). PIE-1::DEGRON::GFP was completely depleted with auxin treatment in (B).

Figure S5 Scatter plot comparing mRNA reads between repeats of each sample

Table S1 PIE-1 interactors identified in Yeast two-hybrid screen

Table S2 List of enriched SUMO-conjugated proteins in *C. elegans*. Mass spectrometry of SUMO purification eluates from nickel affinity chromatography enriched SUMO and SUMO-conjugated proteins in *C. elegans*.

Table S3 Strains and alleles used in this study

Table S4 sgRNA sequences for CRISPR

Table S5 RNA-seq data from dissected gonads of wild-type, *pie-1(ne4303[K68R])*, and *pie-1::degron::gfp*

### Figure S1

## A

| | Lysis B1 | Lysis B2 | Lysis B3 | Lysis B4 | Lysis B5 | Lysis B6 | Lysis B7 | Lysis B8 | Lysis B9 |
|---|---|---|---|---|---|---|---|---|---|
| Hepes 1M pH7.5 | 20µl | 20µl | 20µl | 20µl | 20µl | | | | |
| KAc 1M | 110µl | 110µl | 110µl | | | | | | |
| Tris-Cl 1M pH8.0 | | | | | | 40µl | 40µl | | |
| NaCl 3M | | | | | | 50µl | 50µl | 100µl | |
| Tween20 1% | 100µl | 100µl | | 100µl | 100µl | | | 100µl | 100µl |
| Triton X-100 10% | 50µl | | 50µl | 50µl | 50µl | | | 50µl | 50µl |
| deoxy-bigCHAP 100mM | | | | | | 100µl | 100µl | | |
| Na3 Citrate 1M | | | | 125µl | 250µl | 125µl | 250µl | | |
| NH4-Ac 3M | | | | | | | | 67µl | 500µl |
| MgCl2 1M | 2µl | 2µl | 2µl | 2µl | 2µl | 2µl | 2µl | 2µl | 2µl |
| DTT 1M | 1µl | 1µl | 1µl | 1µl | 1µl | 1µl | 1µl | 1µl | 1µl |
| P solution 100x | 10µl | 10µl | 10µl | 10µl | 10µl | 10µl | 10µl | 10µl | 10µl |
| Protease Inhibitors | √ | √ | √ | √ | √ | √ | √ | √ | √ |

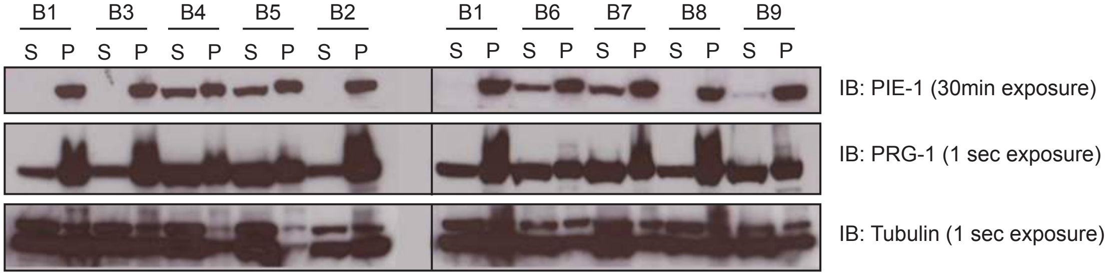


## B

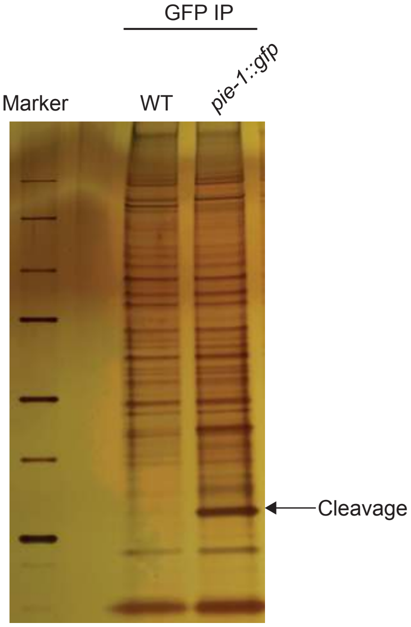


(Silver staining)

## C

S327: Cleavage site

```
MAQTKPIAEQMAALNNSDDTSFAADRSNSLLNATCPARIQNSVD
QRKINRSFNDSLSSGYSGKWLRPKREALKITPLAQIDEAPATKR
HSSAKDKHTEYKTRLCDAFRREGYCPYNDNCTYAHGQDELRVPR
RRQEYYSRDPPRERRDSRSRRDDVDTTINRSSSSASKHHDENRR
PSNNHGSSNRRQICHNFERGNCRYGPRCRFIHVEQMQHFNANAT
VYAPPSSDCPPPIAYYHHHPQHQQQFLPFPMPYFLAPPPQAQQG
APFPVQYIPQQHDLMNSQPMYAPMAPTYYYQPINSNGMPMMDVT
IDPNATGGAFEVFPDGFFSQPPPTIISASSKGEELFTGVVPILV
ELDGDVNGHKFSVSGEGEGDATYGKLTLKFICTTGKLPVPWPTL
VTTFCYGVQCFSRYPDHMKRHDFFKSAMPEGYVQERTIFFKDDG
NYKTRAEVKFEGDTLVNRIELKGIDFKEDGNILGHKLEYNYNSH
NVYIMADKQKNGIKVNFKIRHNIEDGSVQLADHYQQNTPIGDGP
VLLPDNHYLSTQSALSKDPNEKRDHMVLLEFVTAAGITHGMDEL
YKAS*
```

Yellow: PIE-1
Green: GFP

### Figure S2

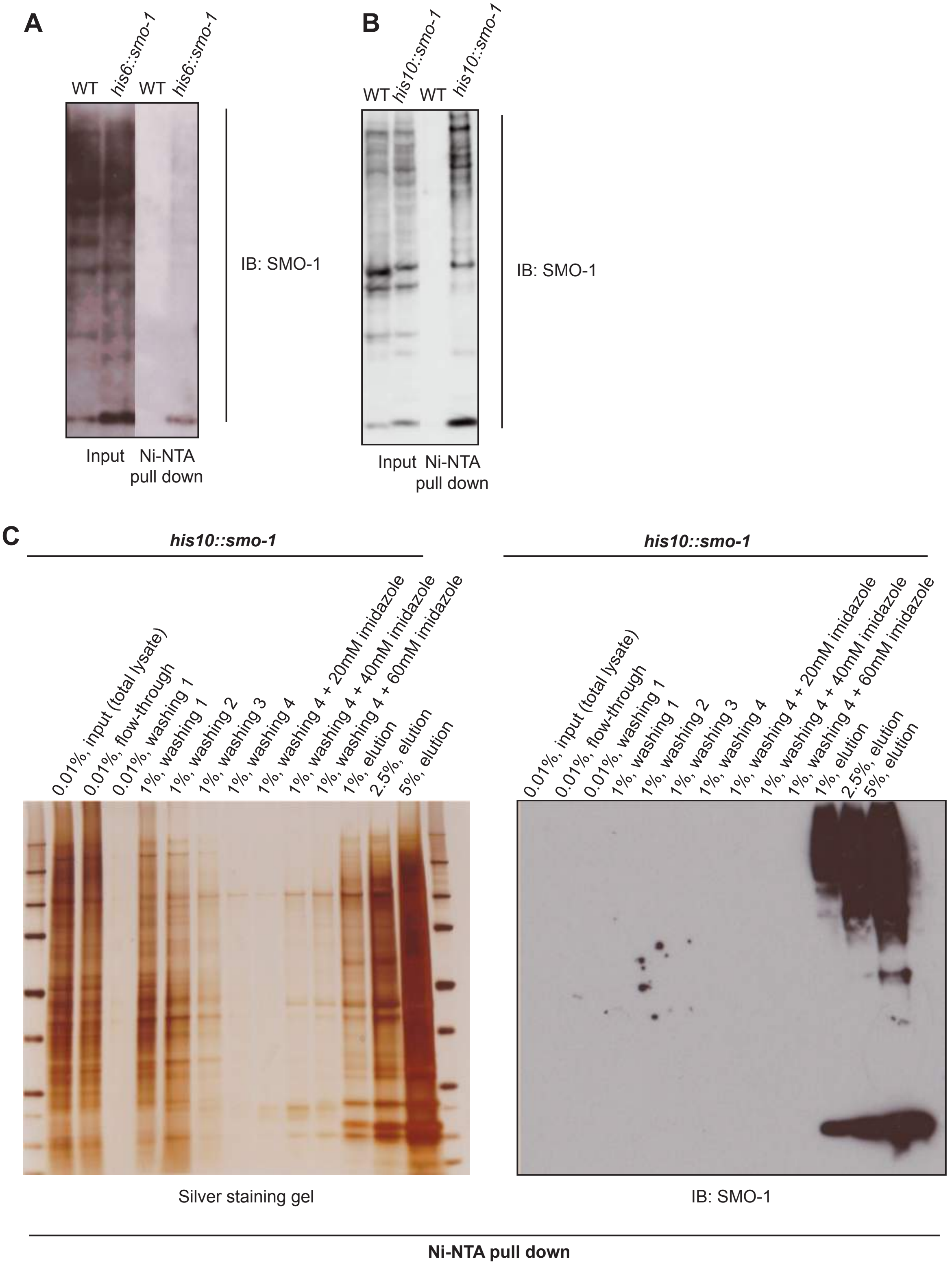
A
WT
his6::smo-1
WT
his6::smo-1
IB: SMO-1
Input
Ni-NTA pull down
B
WT
his10::smo-1
WT
his10::smo-1
IB: SMO-1
Input
Ni-NTA pull down
C
his10::smo-1
his10::smo-1
0.01%, input (total lysate)
0.01%, flow-through
0.01%, washing 1
1%, washing 1
1%, washing 2
1%, washing 3
1%, washing 4
1%, washing 4 + 20mM imidazole
1%, washing 4 + 40mM imidazole
1%, washing 4 + 60mM imidazole
1%, elution
2.5%, elution
5%, elution
Silver staining gel
IB: SMO-1
Ni-NTA pull down

#### Figure S3

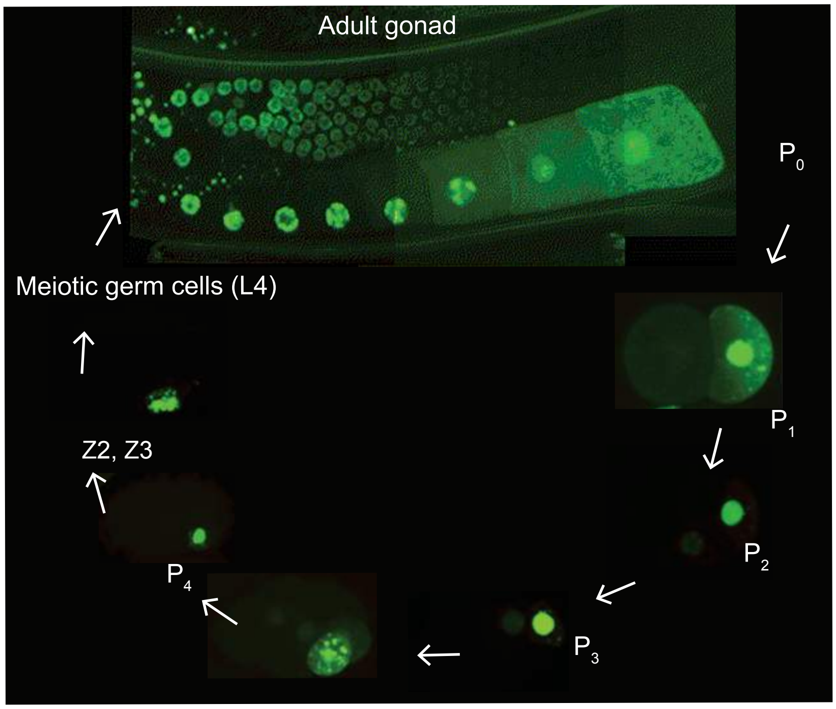

#### Figure S4

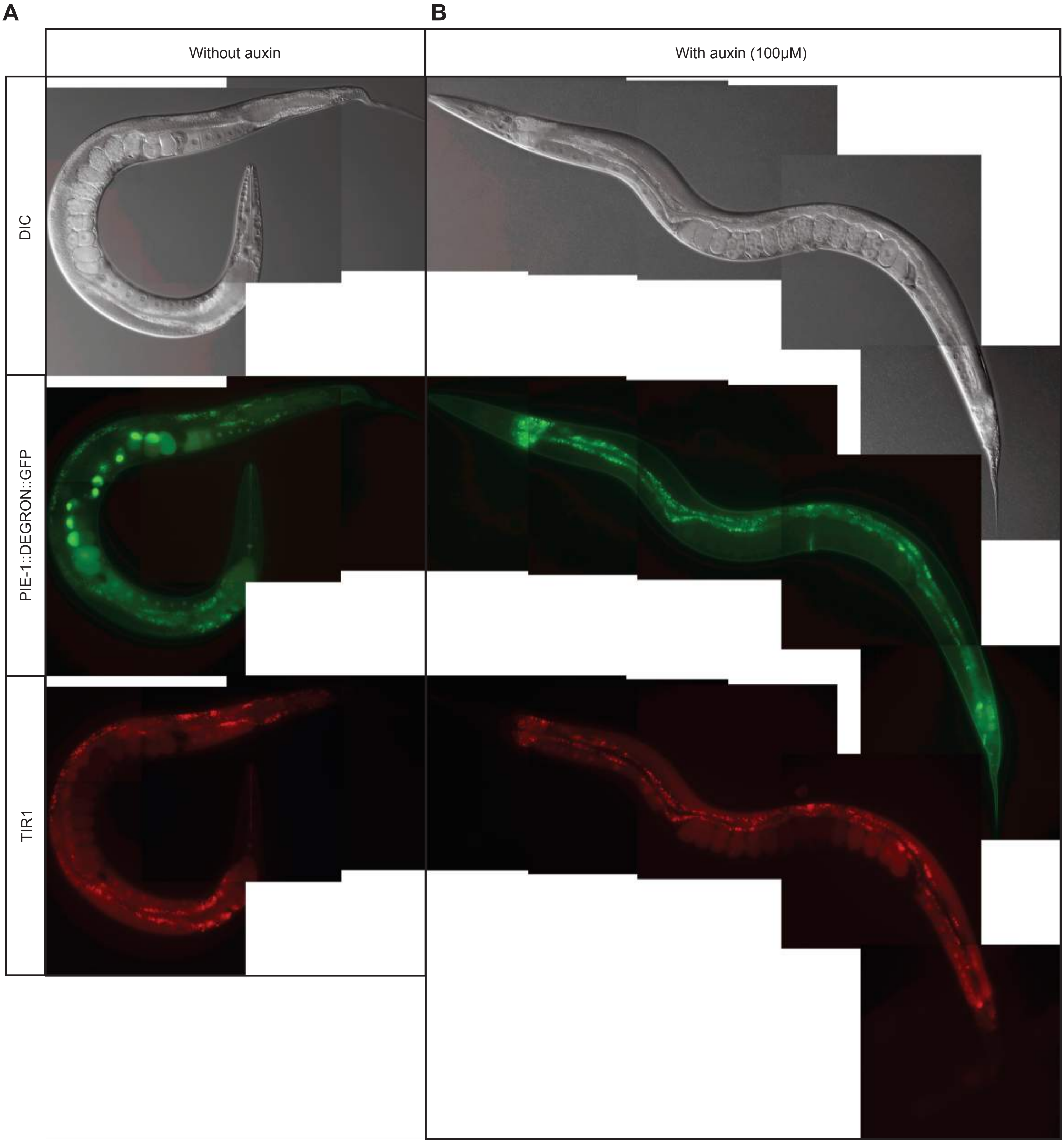

Figure S5

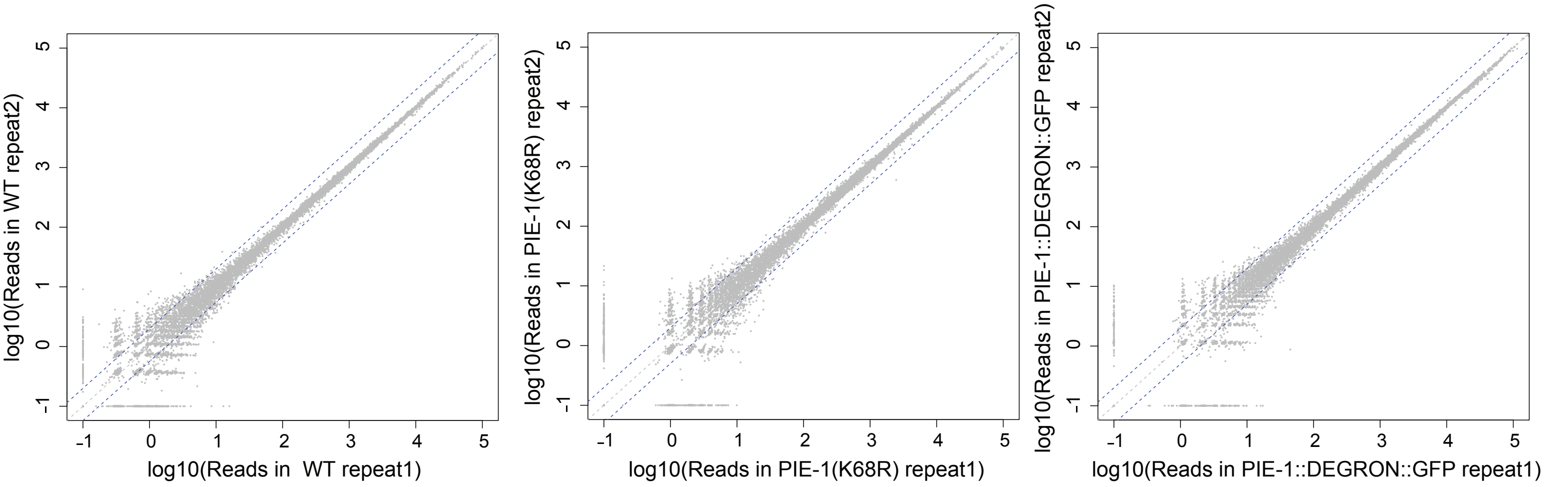
log10(Reads in WT repeat2)
log10(Reads in WT repeat1)
log10(Reads in PIE-1(K68R) repeat2)
log10(Reads in PIE-1(K68R) repeat1)
log10(Reads in PIE-1::DEGRON::GFP repeat2)
log10(Reads in PIE-1::DEGRON::GFP repeat1)
