## Supplemental Table 1 for "PIE-1 promotes SUMOylation and activation of HDAC1 during the *C. elegans* oogenesis"

Table S1. PIE-1 interactors identified in Yeast two-hybrid screen. The score that was computed to assess interaction reliability represents the probability of non-specific interaction. A is the lowest probability and E is the highest probability.

| Gene | Domain/function | Score |
| --- | --- | --- |
| ZIF-1 | ZF interacting protein/ubiquitination | A |
| GEI-17 | E3 ligase for sumo | A |
| GLD-3 | KH domain/Germ line development | B |
| MBL-1 | MUSCLEBLIND-mRNA splicing regulator | B |
| PIE-1 | Probable homodimer | B |
| POP-1 | TCF family member/posterial pharynx defect | B |
| LON-1 | PR-protein superfamily/ a target of TGF beta family | B |
| SMO-1 | SUMO | C |
| UBC-9 | E2 ligase for sumo | E |
| MEP-1 | C2H2/function with NuRD complex | E |
| RAD-26 | An ortholog of human RAD54L2 | E |
|  | A transcription-coupled repair factor |  |
