## Supplemental Table 3 for "PIE-1 promotes SUMOylation and activation of HDAC1 during the *C. elegans* oogenesis"

Table S3 Strains and alleles used in this study

| Strain name: Genotype | Source | Method |
| --- | --- | --- |
| *C. elegans*: Strain JJ532: *pie-1(zu154) unc-25(e156)/qc-1* | *Caenorhabditis* Genetics Center | N/A |
| *C. elegans*: Strain OD56: *unc-119(ed3)*III; Itls37[(pAA64)*pie-1p::mCherry::his-58*+ *unc-119(+)*]IV | (McNally et al., 2016) | MosSCI |
| *C. elegans*: Strain CA1199: *unc-119(ed3)*III; ieSi38[*sun-1p::TIR1::mRuby::sun-1 3’UTR, cb-unc-119(+)*]IV | (Zhang et al., 2015) | MosSCI |
| *C. elegans*: Strain WM330: *pie-1(ne4301*[PIE-1::GFP]*)*III | (Kim et al., 2014) | CRISPR |
| *C. elegans*: Strain WM331: *pie-1(ne4302*[PIE-1::3xFLAG]*)*III | (Kim et al., 2014) | CRISPR |
| *C. elegans*: Strain WM332: *pie-1(ne4303*[PIE-1(K68R)]*)*III | (Kim et al., 2014) | CRISPR |
| *C. elegans*: Strain WM333: *pie-1(ne4304*[PIE-1(K68R)::3xFLAG]*)*III | This study | CRISPR |
| *C. elegans*: Strain WM338: *smo-1(ne4311*[6xHIS::SMO-1]*)*I | This study | CRISPR |
| *C. elegans*: Strain WM650: *pie-1(ne4301*[PIE-1::GFP]*)*III; Itls37[(pAA64)*pie-1p::mCherry::his-58*+ *unc-119(+)*]IV | This study | Cross |
| *C. elegans*: Strain WM651: *ubc-9(ne4446*[UBC-9(G56R)]*)*IV | This study | CRISPR |
| *C. elegans*: Strain WM667: *smo-1(ne4346*[10xHIS::SMO-1]*)*I | This study | CRISPR |
| *C. elegans*: Strain WM671: *mep-1(ne4380*[MEP-1::GFP::TEV::3xFLAG]*)*IV | This study | CRISPR |
| *C.elegans*: Strain WM709: *pie-1(ne4443*[PIE-1::Degron::GFP]*), unc-119(ed3)*III; ieSi38[*sun-1p::TIR1::mRuby::sun-1 3’UTR, cb-unc-119(+)*]IV | This study | CRISPR |
| *C. elegans*: Strain WM716: *pie-1(zu154) unc-25(e156)/qc-1*III*; ubc-9(ne4446*[UBC-9(G56R)]*)*IV | This study | Cross |
| *C. elegans*: Strain WM717: *smo-1(ne4346*[10xHIS::SMO-1]*)*I; *pie-1(ne4302*[PIE-1::3xFLAG]*)*III | This study | Cross |
| *C. elegans*: Strain WM718: *smo-1(ne4346*[10xHIS::SMO-1]*)*I; *pie-1(ne4304*[PIE-1(K68R)::3xFLAG]*)*III | This study | Cross |
| *C. elegans*: Strain WM719: *pie-1(ne4302*[PIE-1::3xFLAG]*)*III; *mep-1(ne4380*[MEP-1::GFP::TEV::3xFLAG]*)*IV | This study | Cross |
| *C. elegans*: Strain WM720: *pie-1(ne4304*[PIE-1(K68R)::3xFLAG]*)*III; *mep-1(ne4380*[MEP-1::GFP::TEV::3xFLAG]*)*IV | This study | Cross |
| *C. elegans*: Strain WM721: *smo-1(ne4346*[10xHIS::SMO-1]*)*I; *mep-1(ne4380*[MEP-1::GFP::TEV::3xFLAG]*)*IV | This study | Cross |
| *C. elegans*: Strain WM722: *smo-1(ne4346*[10xHIS::SMO-1]*)*I; *pie-1(ne4303*[PIE-1(K68R)]*)*III*; mep-1(ne4380*[MEP-1::GFP::TEV::3xFLAG]*)*IV | This study | Cross |
