## Supplemental table 4 for "PIE-1 promotes SUMOylation and activation of HDAC1 during the *C. elegans* oogenesis"

Table S4 sgRNA sequences for CRISPR

| Target gene/ sgRNA sequence | Source |
| --- | --- |
| *pie-1* sgRNA (for c-terminal tagging)  5’-GCTGAGAGAAGAATCCATCG-3’ | (Kim et al., 2014) |
| *smo-1* sgRNA (for N-terminal tagging)  5’-GAGACTCCCGCTATAAACGA-3’ | This study |
| *ubc-9* sgRNA (for G56R)  5’-GGCTCGAACTTGCACTTTGG-3’ | This study |
| *mep-1* sgRNA (for C-terminal tagging)  5’-GCGCAAAAGAAGGAAGACGG-3’ | This study |
